# Language models enable zero-shot prediction of the effects of mutations on protein function

**DOI:** 10.1101/2021.07.09.450648

**Authors:** Joshua Meier, Roshan Rao, Robert Verkuil, Jason Liu, Tom Sercu, Alexander Rives

## Abstract

Modeling the effect of sequence variation on function is a fundamental problem for understanding and designing proteins. Since evolution encodes information about function into patterns in protein sequences, unsupervised models of variant effects can be learned from sequence data. The approach to date has been to fit a model to a family of related sequences. The conventional setting is limited, since a new model must be trained for each prediction task. We show that using only zero-shot inference, without any supervision from experimental data or additional training, protein language models capture the functional effects of sequence variation, performing at state-of-the-art.

## 1 Introduction

Proteins have a myriad of diverse functions that underlie the complexity of life. Protein sequences encode function via structure through the spontaneous folding of the sequence into the three dimensional structure of the protein [1]. The effects of sequence mutations on function form a landscape that reveals how function constrains sequence. Alterations at some sites in a protein sequence cannot be tolerated because they are essential to the protein’s function. Other sites evolve together because the structure and function is determined by them collectively. Mutations can enhance the activity of a protein, attenuate it, or leave it unchanged.

The functional effect of sequence variations can be measured through deep mutational scanning experiments [2]. Consisting of thousands to hundreds of thousands of measurements of protein function, deep mutational scans give insight into the intrinsic constraints on a protein’s structure and function. Due to the cost and difficulty of implementing such experiments, compilations of deep mutational scanning data include experiments on a few dozens of proteins at most, relative to the tens of thousands of proteins encoded in the human genome, and the millions more across the tree of life that we would like to understand.

A model that learns the landscape linking sequence to function can provide insight into function without having to do experiments. Unsupervised models of mutational effects can be learned from sequences [3, 4]. Statistical patterns in a family of evolutionarily related protein sequences contain information about structure and function [5–7]. This is because the properties of a protein act as constraints on the selection of sequences through evolution [8].

In the natural language modeling community, there has been interest in zero-shot transfer of models to new tasks. Massive language models can solve tasks they haven’t been directly trained on [9–11]. Recently protein language models have achieved state-of-the-art in various structure prediction tasks [12–14]. Work to date has mainly focused on transfer in the classical representation learning setting, using pre-trained features with supervision on the downstream task.

In this work we show that language models trained on large and diverse protein sequence databases can predict experimental measurements of protein function without further supervision. Prior work has focused on transferring the representations using supervision from experimental data [15, 16]. We find that language models can transfer to predict functional measurements without supervision. Language models perform *zero-shot* and *few-shot* prediction of mutational effects across a variety of proteins with widely differing functions. We perform experiments with state-of-the-art protein language models ESM-1b [12] and MSA Transformer [13]. We introduce a new protein language model, ESM-1v, with zero-shot performance comparable to state-of-the-art mutational effect predictors. Performance can be further improved by fine-tuning the model with sequences from the protein family. Predictions capture the functional landscape of the protein, correlate with amino acid conservation patterns in the core and surface, and identify residues responsible for binding and activity.

## 2 Zero-shot transfer

Zero-shot learning has classically described the extension of a classifier to a new set of classes that have not been seen in training [17]. In natural language processing this idea has been extended to describe the transfer of models to entirely new tasks without further training. Proposed as zero-data learning by Larochelle et al. [18], this perspective on transfer has been at the center of recent work understanding the generalization capabilities of large language models [9–11, 19]. The distinction from representation learning is that the models are used *directly* without additional supervision for the task. This means that the tasks must be learned purely from pre-training.

In this work we take a similar perspective on zero-shot transfer to that of GPT-3, described in Brown et al. [10]. We define zero-shot transfer to be transfer of a model to a new task without any further supervision to specialize the model to the task. We also consider the closely related idea of few-shot transfer. Here as in Brown et al. [10] we define the few-shot setting to be one in which a few positive examples are given to the model as inputs at inference time. As in the zero-shot setting, no gradient updates are performed to specialize the model. Similar to Brown et al. [10], the claim is not one of out-of-distribution generalization. The assumption is that in the pre-training stage, the model learns information relevant to the tasks to which it will later be transferred. In the case of protein language models, the pre-training dataset includes sequences from across evolution, which implies the model may see examples of sequences from protein families on which it will be evaluated. The essential departure from the standard approach in computational biology is that the model is *general purpose* and can be applied across a variety of tasks without specialization.

Measurements of function, a property of central importance to the understanding and design of proteins, are a practical ground for studying the generalization capability of protein language models. Deep mutational scanning experiments measure the effects of thousands to hundreds of thousands of mutations on a single protein, and have been performed on a variety of proteins having different functions and using various forms of experimental measurement. We study zero-shot and few-shot transfer of protein language models to function prediction using this data.

Supervised methods trained with data from experimental measurements [15, 16], and unsupervised methods trained only on sequences [3, 4] have been developed for prediction of mutational effects. Unsupervised mutational effect predictors are trained as task specific models on sequences from an individual protein family. In this view every protein is an independent prediction task where the objective is to score the effect of mutations on the protein’s function. While mutational effect predictors trained on multiple sequence alignments (MSAs) are typically described as unsupervised, they can also be seen as weakly supervised. Hsu et al. [15] observe that such models have weak supervision on the task through the MSA, which describes the fitness landscape of the protein through positive examples.

If protein language models can learn the information necessary to solve a task from pre-training, then they can be applied *directly* to new instances of the task, without specialization. This would mean that in practice a single general purpose model can be trained once and then applied to a variety of possible tasks. Thus zero-shot and few-shot transfer represent fundamentally new unsupervised learning capabilities that protein language models can bring to the computational biology toolkit.

## 3 Method

Protein language models trained with the masked language modeling objective are supervised to output the probability that an amino acid occurs at a position in a protein given the surrounding context. We use this capability to score sequence variations. For a given mutation we can consider the amino acid in the wildtype protein as a reference state, comparing the probability assigned to the mutated amino acid with the probability assigned to the wildtype.

We score mutations using the log odds ratio at the mutated position, assuming an additive model when multiple mutations *T* exist in the same sequence:

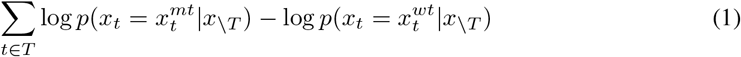

Here the sum is over the mutated positions, and the sequence input to the model is masked at every mutated position.

### 3.1 Zero-shot and few-shot transfer

In the zero-shot setting, inference is performed directly on the sequence to be evaluated. Since the MSA Transformer can take multiple sequences as input at inference time, we use this model in the few-shot setting, where additional sequences from the protein family are provided along with the sequence to be evaluated. In both the zero-shot and few-shot settings, only forward passes of the models are performed during inference; no gradient updates are taken. Fig. 2 illustrates the approach in comparison to the current practice of fitting a new model for each task.

**Figure 1:**
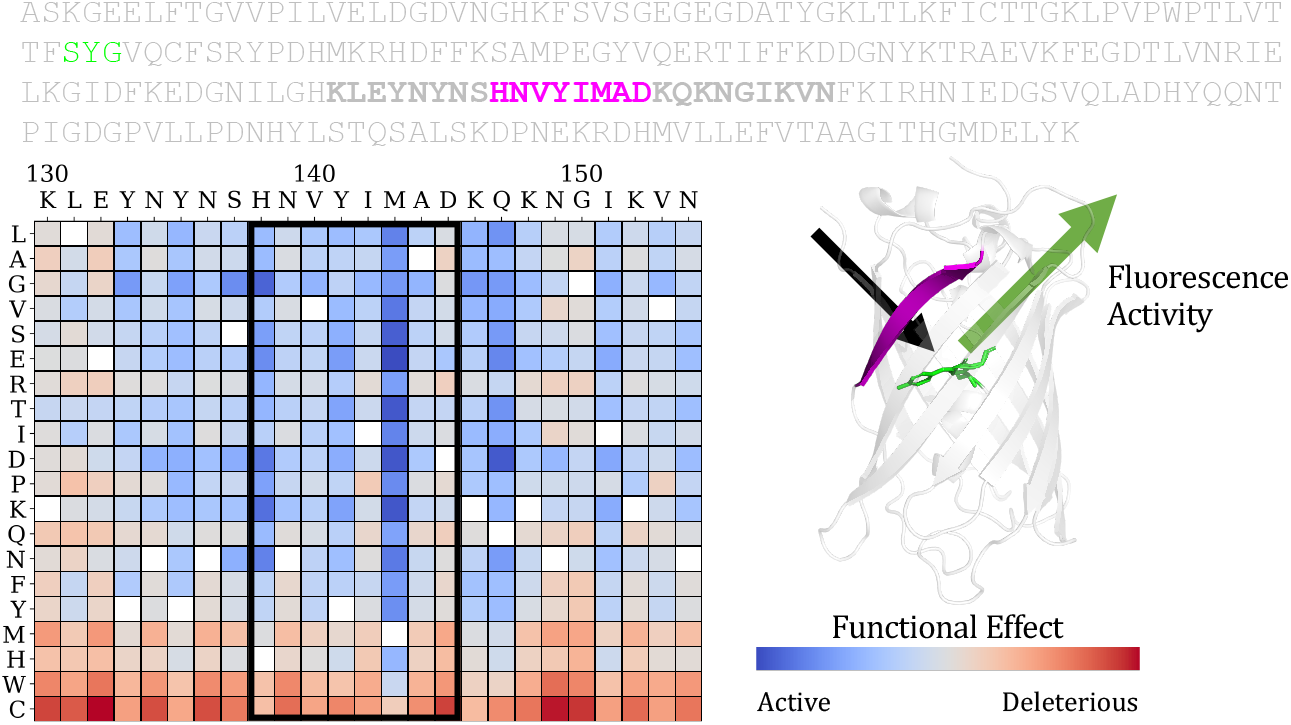
Depiction of a mutational effect prediction task. The objective is to score the effect of sequence mutations on the function of a protein. Deep mutational scanning experiments provide ground truth experimental measurements of the protein’s function (fluorescence activity in the example here) for a large set of single mutations or combinations of mutations. For each protein, the prediction task is to score each possible mutation and rank its relative activity. Predictions for single substitutions can be described in a score matrix. The columns are the positions in the sequence. The rows are the possible variations at each position.

**Figure 2:**
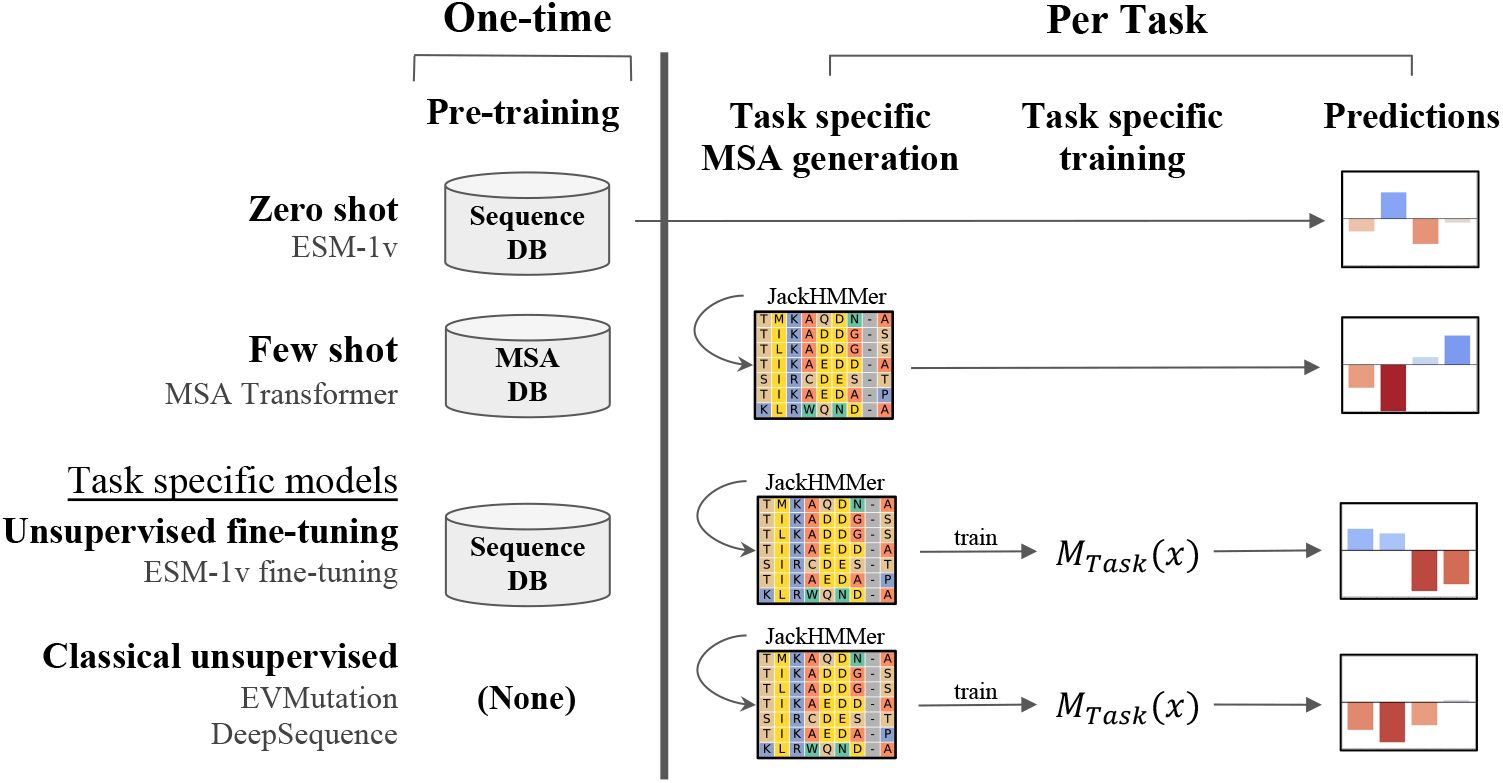
Steps involved in variant effect prediction methods. Compared with EVMutation [4] and DeepSequence [20], MSA Transformer and ESM-1v require no task-specific model training for inference. Moreover, ESM-1v does not require MSA generation.

### 3.2 Inference efficiency

Inference with ESM-1v is more efficient than current state-of-the-art methods. This is a result of two important differences: (i) the effect of mutations can be inferred directly without training a task-specific model; (ii) fitness landscapes can be predicted with a single forward pass. Time requirements are summarized in Fig. 7.

### 3.3 Scoring with MSA Transformer

We score mutations with MSA Transformer using the log odds ratio and additive model in Eq. (1). However, since MSA Transformer uses a set of sequences for inference, we input the sequence to be evaluated as the first sequence, and provide additional sequences from the MSA as context. Masking and scoring are performed on the first sequence only.

## 4 Results

### 4.1 Experimental setup

#### Prediction Models

We compare to state-of-the-art unsupervised variant prediction methods, EVMutation [4] and DeepSequence [20]. We also examine performance of a variety of protein language models that have been recently introduced in the literature.

The position specific scoring matrix (PSSM), EVmutation [4], and DeepSequence [20] methods are all MSA based. The PSSM treats each position in the sequence independently, factorizing the likelihood into one term per sequence position. EVmutation is a Potts model, which adds pairwise terms modeling the interactions between positions. DeepSequence introduces a latent code, allowing potential higher-order interactions between positions.

UniRep [21], TAPE [22], ProtBERT-BFD [14], ESM-1b [12], and ESM-1v (introduced here), are all single-sequence language models trained on large databases of unaligned and unrelated protein sequences (e.g. Pfam [23] or UniRef [24]). With the exception of UniRep, which is trained using next token prediction, all models are trained with masked language modeling [25].

Finally, the MSA Transformer [13] is a combination of both approaches; it is trained on a large database of MSAs using masked language modeling and takes an MSA as input during inference.

#### ESM-1v

We train ESM-1v, a 650M parameter transformer language model for prediction of variant effects, on 98 million diverse protein sequences across evolution. The model is trained only on sequences, without any supervision from experimental measurements of function. We use Uniref90 `2020-03` [24], employing the ESM-1b architecture and masked language modeling approach of Rives et al. [12]. The model attains a perplexity of 7.29 on a set of held-out Uniref90 sequences (Table 10). We train five models with different seeds to produce an ensemble.

#### Evaluation

Models are evaluated on a set of 41 deep mutational scans collected by Riesselman et al. [20], which comprise a variety of tasks assessing a diverse set of proteins. Across tasks, the experiments differ in the functions tested and in the measurements performed. We treat each deep mutational scanning dataset as a separate prediction task, scoring each of the variants in the dataset with the model. The tasks are split into a validation set of ten mutational scanning datasets and a test set consisting of the remaining datasets. We evaluate performance by comparing the scores with the experimental measurements using Spearman rank correlation.

#### Comparisons

Since the published versions of EVMutation and DeepSequence use MSAs generated from an earlier version of Uniref100, we generate new MSAs using EVMutation methodology and the version of Uniref100 concurrent with our pretraining dataset. We train replications of EVMutation and DeepSequence using their open source code. The same MSAs are also used in few-shot experiments with MSA Transformer and unsupervised fine-tuning experiments with ESM-1v.

### 4.2 Language models enable zero-shot and few-shot prediction of the effects of mutations

ESM-1v and MSA Transformer models make state-of-the-art predictions. Table 1 compares overall performance of the models across the 41 mutational scanning datasets. Fig. 3 presents a comparison between ESM-1v and DeepSequence on each of the tasks. Zero-shot inference with ESM-1v has a better correlation with experimental measurements than DeepSequence on 17 of the 41 datasets. The two methods are not statistically distinguishable via a paired *t*-test.

**Table 1:** Comparison of protein language models to state-of-the-art methods. Average |Spearman *ρ*| on full and test sets. DeepSequence and ESM-1v models are each ensembles of 5 models. MSA Transformer is a single model, but is ensembled across 5 random samples of the MSA.

| Models | Full | Test |
| --- | --- | --- |
| PSSM | 0.460 | 0.460 |
| EVMutation (published) | 0.508 | 0.495 |
| EVMutation (replicated) | 0.511 | 0.498 |
| DeepSequence (published) | 0.514 | 0.499 |
| DeepSequence (replicated) | 0.520 | 0.506 |
| MSA Transformer | 0.542 | 0.524 |
| ESM-1v (zero shot) | 0.509 | 0.482 |
| ESM-1v (+further training) | 0.538 | 0.519 |

**Figure 3:**
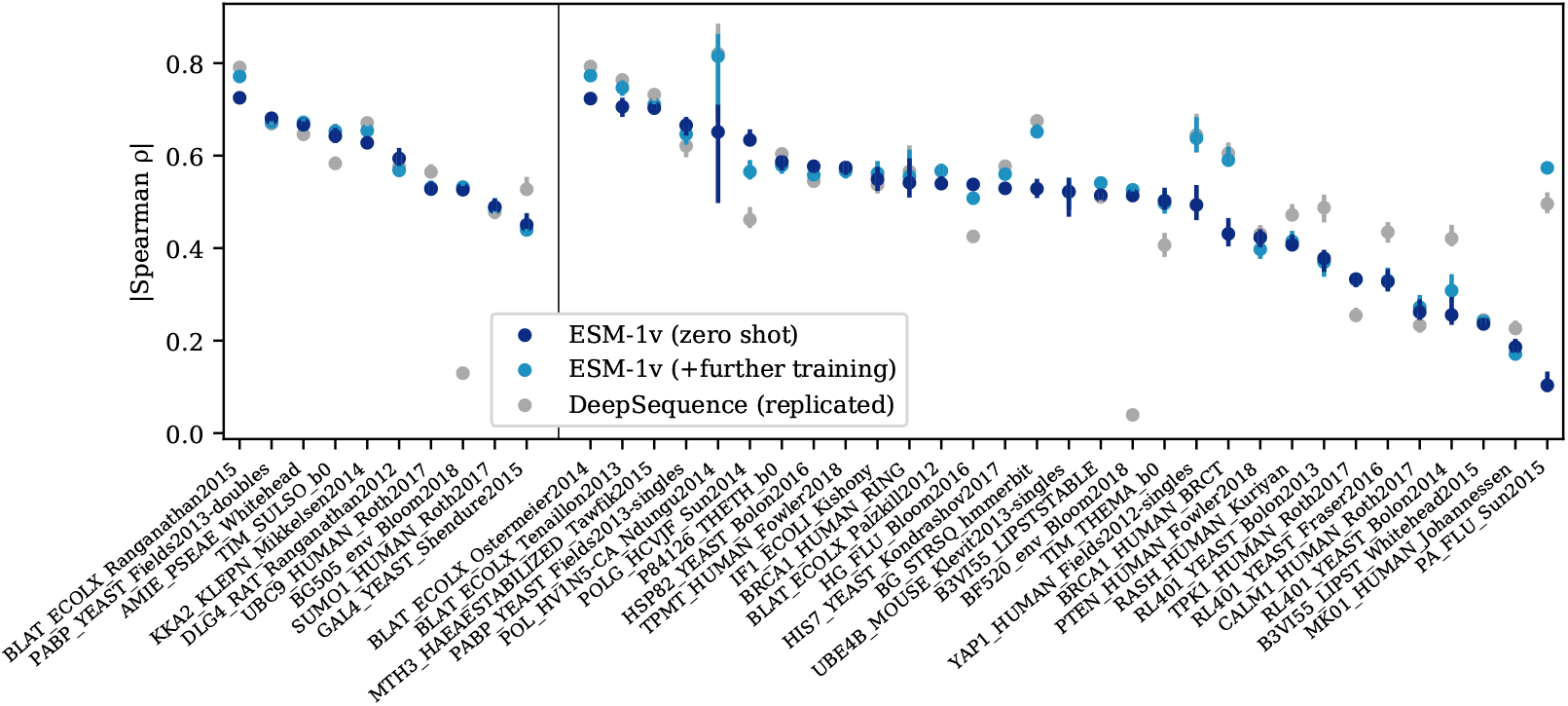
Per task performance. Comparison across 41 deep mutational scanning datasets. Points are |Spearman *ρ*| on each dataset, error bars show standard deviation of 20 bootstrapped samples. Validation proteins are shown to the left of the dividing line and test proteins to the right. In 17 out of the 41 tasks, ESM-1v zero-shot has a higher |Spearman *ρ*| than DeepSequence.

Table 2 compares protein language models in the zero-shot setting. ESM-1v outperforms existing protein language models TAPE [22], UniRep [21], ProtBERT-BFD [14], and ESM-1b [12]. Fig. 8 breaks down performance across each of the tasks.

**Table 2:** Zero-shot performance. Average |Spearman *ρ*| on full and test sets. ^†^Average performance of five ESM-1v models. ^*^Ensemble of the five ESM-1v models.

| Models | Full | Test |
| --- | --- | --- |
| UniRep | 0.156 | 0.151 |
| TAPE | 0.171 | 0.175 |
| ProtBERT-BFD | 0.428 | 0.399 |
| ESM-1b | 0.459 | 0.424 |
| ESM-1v<sup>†</sup> | 0.484 | 0.457 |
| ESM-1v<sup>*</sup> | 0.509 | 0.482 |

#### Pre-training data

We examine the effect of the clustering level of pre-training data. Fig. 4 compares models pre-trained on datasets clustered at increasing sequence identity thresholds. ESM-1b is trained on sequences clustered at a 50% identity threshold. Improvements are seen using a 70% threshold with greatest improvement at 90%. Uniref100 performance appears to deteriorate early in training despite being the largest of the datasets. These results establish a link between model performance and the data distribution, highlighting the importance of training data in the design of protein language models.

**Figure 4:**
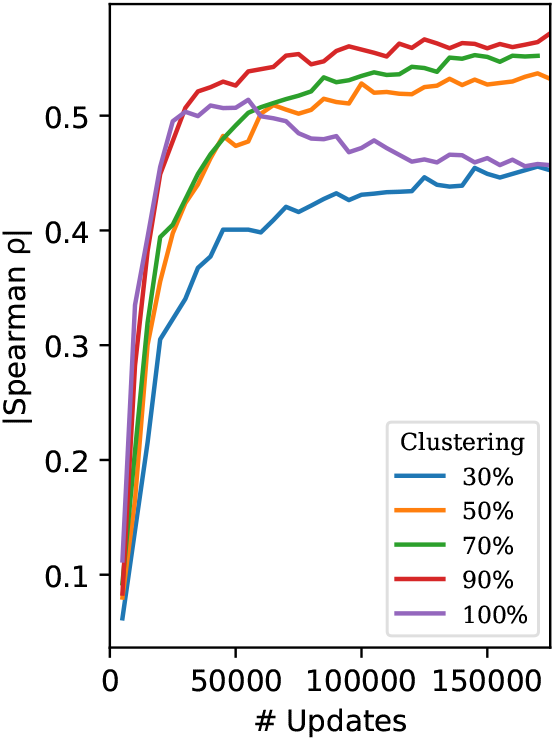
Comparison of pre-training datasets. Average |Spearman *ρ*| on the single-mutation validation set. While a 50% clustering threshold was used for ESM-1b, training with 90% clustering results in a significant improvement on variant prediction tasks. Notably, models trained on Uniref100, the largest dataset in this figure, appear to deteriorate early in training. These results establish a link between model performance and the data distribution, and highlight the importance of training data in the design of protein language models.

#### Scoring methods

We compare four scoring methods on the validation set - masked marginals, wildtype marginals, mutant marginals, and psuedolikelihood. Table 5 shows that the masked marginal approach described in Eq. (1) outperforms other scoring methods, including ones in which the likelihood changes at non-mutated positions are considered. The scoring methods are described in detail in Appendix A.

#### Parameter count

Previous work with protein language models has established a link between model scale and learning of protein structure [12, 26]. We examine zero-shot transfer performance as a function of parameter count. We train models using the same width, depth, and learning rate as described in Henighan et al. [27], observing improvements with scale (Fig. 9). These findings suggest that continued scaling of the models will further improve results.

### 4.3 MSA Transformer

We examine how the sequences provided to MSA Transformer affect few-shot transfer. Table 8 compares sequence selection methods that vary the diversity of the sequences. Providing a more diverse set of sequences improves few-shot performance. Selecting a set of sequences to maximize diversity outperforms selecting a diversity minimizing set of sequences. Random sampling performs even better, and sampling sequences according to sequence weights [28] performs best.

We also vary the number of sequences used for inference. Fig. 11 shows few-shot performance as a function of the number of sequences given as input. The model performs well using only a few sequences, but performs best with 384 total sequences. In the main tables we report results sampling 384 sequences using sequence reweighting and ensembling predictions over five different subsamples from the MSA.

### 4.4 Unsupervised fine-tuning on MSAs

While ESM-1v performs well when evaluated in the zero-shot setting, we explore whether results can be improved by fine-tuning on the MSA. Fine-tuning on MSAs has been used in previous work [21, 16] as a stage in transfer learning to specialize a pre-trained model to a protein family, before applying supervision with labeled data. Here we consider using the fine-tuned model to make unsupervised predictions directly, without adding supervision from experimental data.

We observe that naively fine-tuning the model on the MSA results in rapid overfitting and poor performance on the prediction tasks (Fig. 12). While we experiment with a variety of approaches to freezing parameters during fine-tuning, detailed in Appendix B, none produce significant improvements. We find that an approach using pre-training sequences to regularize the fine-tuning performs well and enables training of all parameters without overfitting (Fig. 13). Spiked fine-tuning improves average absolute Spearman rho on the full dataset from 0.510 for zero-shot evaluation to 0.537 with fine-tuning.

## 5 Analysis of models

### Protein structure and function

ESM-1v probabilities reflect the functional properties of sites within the protein. We use the entropy of the model’s predictions for a position as a measure of its estimation of conservation. The lowest entropy predictions cluster at binding sites. Fig. 14 compares the distribution of the model’s entropy between binding sites and non-binding sites. A significant difference is observed between the entropy assignment to binding and non-binding site residues. Fig. 5 visualizes the side chains of the 10 lowest entropy residues as predicted by the model on the crystal structure of DNA methyltransferase M.HaeIII interacting with its DNA substrate. In the crystal structure a cytosine of the substrate is inserted into the active site of the enzyme. The low entropy residues cluster in the active site and interact with the cytosine. Additional examples are visualized in Fig. 18.

**Figure 5:**
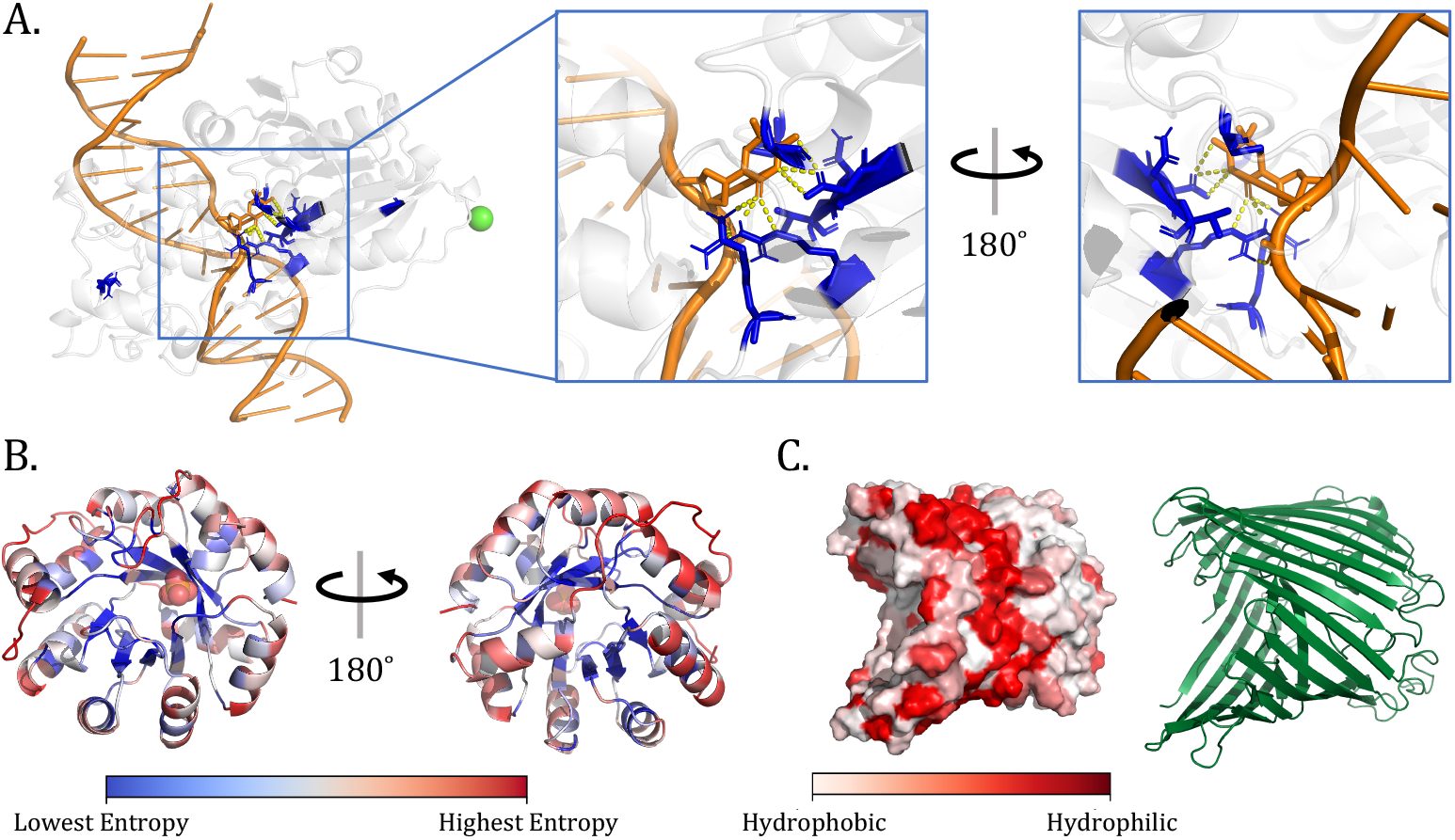
ESM-1v reflects the molecular basis of function in proteins. **(A)** DNA methylase HaeIII (pdbid: 1DCT [29]). Side chains for the top 10 positions with lowest prediction entropy shown in blue. Low-entropy positions cluster in the active site. **(B)** TIM Barrel (pdbid: 1IGS [30]) with residues colored by entropy. The model’s predictions for residues on the surface have highest entropy (red) while those in the core have lower entropy (blue). Notably, residues on the alpha helices show a clear gradient from high to low entropy as residues transition from surface-facing to core-facing. **(C)** Sucrose-specific Porin (pdbid: 1A0T [31]), a transmembrane protein. The model predicts a hydrophobic band where the protein is embedded in the membrane.

The model probabilities also correspond to structure. Fig. 15 compares the entropy assigned to sites that are buried in the core of the protein vs. exposed on the surface. The model assigns significantly lower entropy to sites that are in the core of the protein, consistent with the idea that tight packing in the core places greater constraints on the selection of residues. Fig. 5B visualizes the entropy assigned by the model to each position overlayed on the structure of Indole-3-glycerolphosphate Synthase, a TIM barrel protein. Higher entropy is assigned to residues having outward facing side chains on the alpha helices, while lower entropy is assigned to the inward facing positions. Fig. 17 compares the probability assigned to hydrophobic, polar, and charged amino acids for buried sites vs. non-buried sites. The model prefers hydrophobic residues in the core and hydrophilic residues on the surface. The model probabilities closely match the empirical probabilities and those from the PSSM. Fig. 5C visualizes probability assigned to hydrophobic amino acids on the structure of Sucrose-specific Porin, a transmembrane protein. The model predicts a hydrophobic band in the center where the protein embeds in the membrane.

### Calibration

We evaluate model calibration using 15008 sequences with length < 1024 from the trRosetta [32] dataset. ESM-1v probabilities for each amino acid at each position are calculated with the masked marginal probability in Eq. (1). Fig. 6 shows that the model is generally well calibrated for all amino acids except Methionine. ESM-1v always predicts Methionine as the first position in the sequence since full protein sequences always start with it, so care must be used when applying the model to subsequences. When excepting the first residue, the model achieves an average calibration error (defined for the multi-class setting in Appendix D.4) of 0.006.

**Figure 6:**
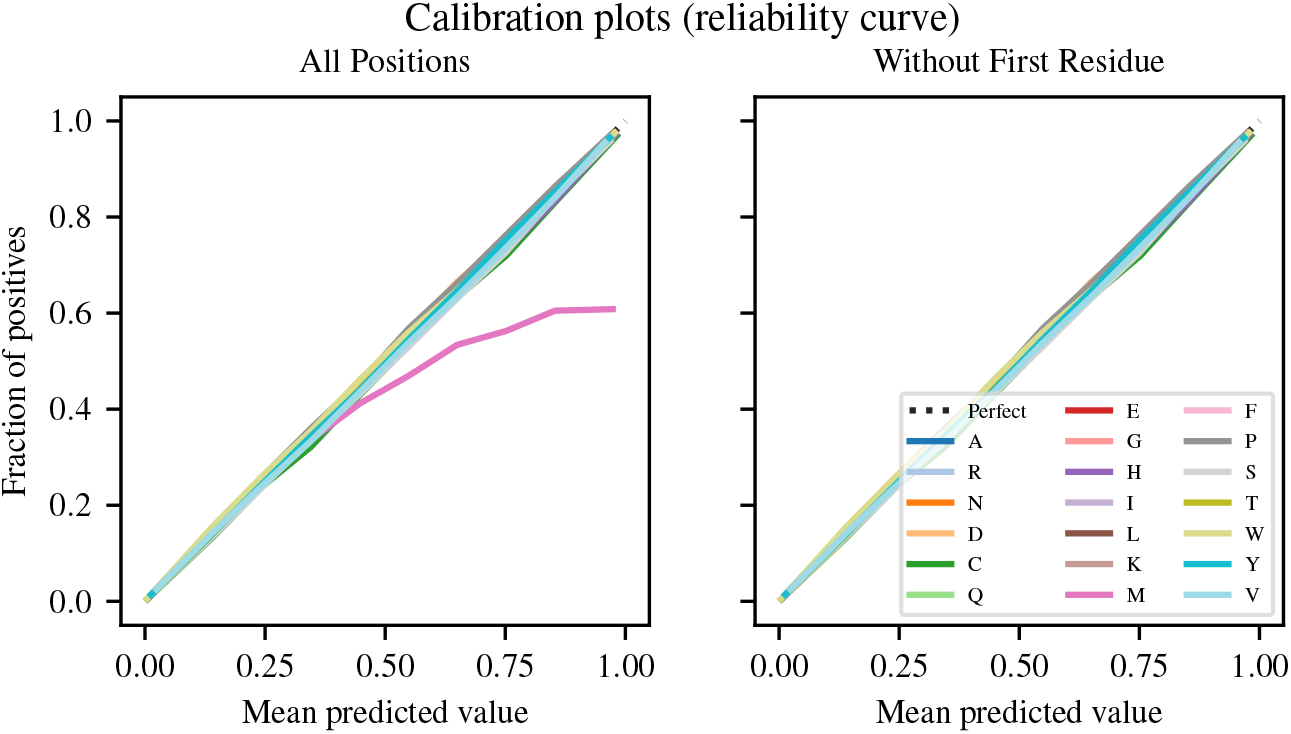
Calibration plot for ESM-1v predictions on each of the 20 naturally occurring amino acids on the trRosetta dataset. The multi-class classification is converted into a set of 20 one-versus-all classifications for the purpose of this analysis. Left and right plots show calibration of all positions and positions excluding the first residue, respectively. Since full sequences always start with Methionine, the model overwhelmingly predicts it in the first position. When evaluating the model on subsequences, such as those in the trRosetta dataset, this causes a miscalibration at the first residue. Including the first residue, the model has an average calibration error (ACE) of 0.011 in the first case and 0.006 in the second.

We also explore the relationship between conservation (entropy of the PSSM) and the model’s predicted entropy. Fig. 16 shows that these are well correlated (Pearson’s *r* = 0.44), suggesting the model is able to identify conserved positions.

## 6 Related Work

### 6.1 Protein language models

In the past few years, a number of groups have developed language models for protein sequences [21, 33, 14, 34, 35, 22, 13, 12]. These models have been used for many tasks, including supervised low-N function prediction [16, 12], remote homology detection [22, 12], and protein generation [35]. The approach to the tasks typically involves transfer learning, where a pretrained language model is fine-tuned for a particular problem. Vig et al. [36] and Rao et al. [26] found that transformer attention corresponds to known biological properties such as structure and binding sites and can be used to predict contacts.

### 6.2 Mutation effect prediction

Supervised and unsupervised methods have been developed for prediction of mutational effects. Supervised methods train models using experimental measurements or labels from databases of clinical variants. Standard machine learning tools including linear regression, random forests, and support-vector machines can be used [37]. Models have been designed specifically for proteins, using feature engineering such as Envision [38] and PolyPhen-2 [39], ensemble methods such as Revel [40], MPC [41], CADD [42], and M-CAP [43], language models such as UniRep [21, 16] and ESM [12], and other representation learning approaches [44, 45].

Unsupervised mutation effect predictors work by inferring the likelihood of a mutation from the evolutionary landscape of the original protein. A density model fit to related sequences is used for scoring. SIFT [46] is a first order approach using a position-specific-scoring-matrix. EVMutation [4] extends this to a second-order approach by training a Potts model on the MSA. DeepSequence [20] includes higher-order interactions by training a VAE on the MSA instead, using the ELBO to score mutations. Riesselman et al. [47] proposes using an autoregressive model that does not require the sequences to be aligned.

Hsu et al. [15] show that unsupervised mutational effect predictors can be extended to perform supervised predictions, with better unsupervised predictors generally resulting in better supervised predictors. This suggests improving unsupervised prediction can drive progress in both settings. Concurrent with our work, Hie et al. [48] use open-source protein language models ESM-1b and TAPE to predict the direction of evolution in protein fitness landscapes.

## 7 Discussion

Advances in language modeling at scale are bringing the goal of a general purpose model for proteins closer to realization. This line of work aspires to a model that learns to read and write biology in its native language, that can be directly applied across a range of protein understanding and design tasks. For scalability, learning from sequences is important: while there are no central databases of high-throughput functional measurements, and few compilations exist, billions of sequences are available to learn from in sequence databases [49, 50]. Sequences give an unparalleled view into the vast diversity and complexity of molecular parts invented by nature through billions of years of evolution.

Unsupervised structure [51–53, 28, 54, 55] and function [3, 4] learning methods first effectively realized the idea that biological properties could be read directly from sequences without supervision from experimental measurements. However these methods are not general purpose in the sense that a specialized model must be trained for every protein for which a prediction is to be made. We show that the same performance can be realized by a general purpose model that has been trained across many diverse protein families. Similar to observations on the learning of tertiary protein structure in large language models [12, 26], we find that increasing the scale of models leads to improvements in function learning. The understanding of mutational landscapes in the models correlates with the molecular basis of function in proteins, capturing binding sites and amino acid preferences that are determined by the folded structure.

Zero-shot transfer is an interesting capability of large scale language models, and represents a major point of departure from the unsupervised learning methods that are the basis for current state-of-the-art inference of protein structure and function. The capability for zero-shot transfer implies that a model can be trained once and then applied to perform inference for many tasks. It is also a window into deeper questions about the forms of generalization that are possible in learning from sequences. Reading structural and functional design principles from sequences is a necessary capability for writing new biologically active sequences. Generalization in the zero-shot setting suggests the potential for large language models to capture knowledge that can be transferred to generating new functional proteins.

## Acknowledgments and Disclosure of Funding

We thank Naman Goyal, Chloe Hsu, Zeming Lin, Ishan Misra, Myle Ott, Sergey Ovchinnikov, and Ethan Perez for valuable input on the paper. Roshan Rao was supported by funding from Facebook, Berkeley Deep Drive, and DARPA XAI.

## A Extraction methods

ESM-1v is pre-trained to output the probability for each possible amino acid at a masked position. We explore four methods of scoring the effects of mutations using the model:

- **Masked marginal**: Probabilities are extracted according to the mask noise during pretraining. At each position, we introduce a mask token and record the model’s predicted probabilities of the tokens at that position.
- **Mutant marginal**: Probabilities are extracted according to the random token noise during pre-training. Among the 15% predicted positions in the sequence during pre-training, 10% of those are randomly mutated and 10% retain their original identities. The model is tasked to predict the correct token at those positions. Therefore, in this extraction method, we follow the pre-training methodology by passing in mutated tokens and recording the model’s probability that they are correct.
- **Wildtype marginal**: We perform a single forward pass using the wildtype sequence. This method enables fast scoring as just a single forward pass is used.
- **Pseudolikelihood**: This method is proposed in the literature for scoring with masked language models [86].

In all cases, we assume an additive model when multiple mutations are present in a sequence. Results are summarized in Tables 5 and 7.

**Table 3:** Zero-shot learning is a natural extension of the various approaches that have been used for mutational effect prediction to date. Rather than training a new model for every task, a single general purpose model is trained and can be directly applied across multiple tasks. The approach is fully unsupervised, no information from experimental measurements of function is used.

| Approach | Formal setting | Task level supervision | Representative Methods |
| --- | --- | --- | --- |
| Supervised mutation prediction | Supervised | Direct supervision from experimental measurements | Reviewed in [87] |
| Model trained on sequences from individual family | Unsupervised | Weak positive supervision from MSA | [4, 20] |
| Fine-tuning on experimental data | Semi-supervised transfer | Supervision from experimental measurements | [21, 16, 12] |
| Fine-tuning on MSA | Transfer learning with weak-positive supervision | Weak positive supervision from MSA | Introduced for transfer learning in [21] |
| Direct forward pass | Zero-shot learning | None | This work |

Let *x^mt^* and *x^wt^* represent the mutant and wildtype sequences. We refer to *x_−i_* as the sequence *x* with a mask introduced at position *i*. We refer to the set of mutations that are introduced as the set *M*. For example, if mutations are introduced at positions 3 and 6, then *M* = {3, 6}.

## Masked marginal probability (L forward passes)

This method performs best among the four. We introduce masks at the mutated positions and compute the score for a mutation by considering its probability relative to the wildtype amino acid (Strategy a):

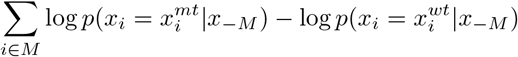

**Figure 7:**
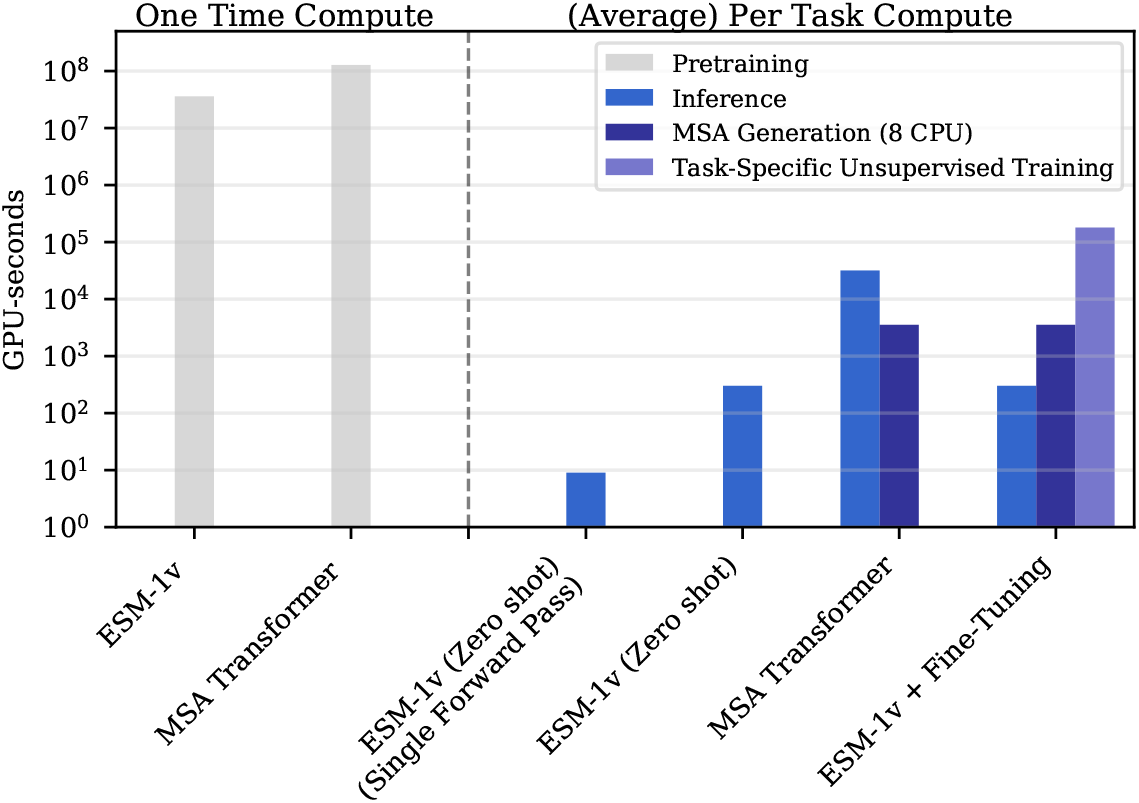
Compute requirements in GPU-seconds for (left) pre-training and (right) average task. With open-sourced pre-trained models, end users bypass the pre-training phase and only incur inference costs. ESM-1v and MSA Transformer amortize compute cost into a single expensive pre-training run. After pre-training, inference is fast. On average, it takes 10 seconds to label a deep mutational scan from Riesselman et al. [20] with ESM-1v (Zero-shot, Single Forward Pass). Performance improves marginally with the more expensive scoring scheme (Table 5).

**Figure 8:**
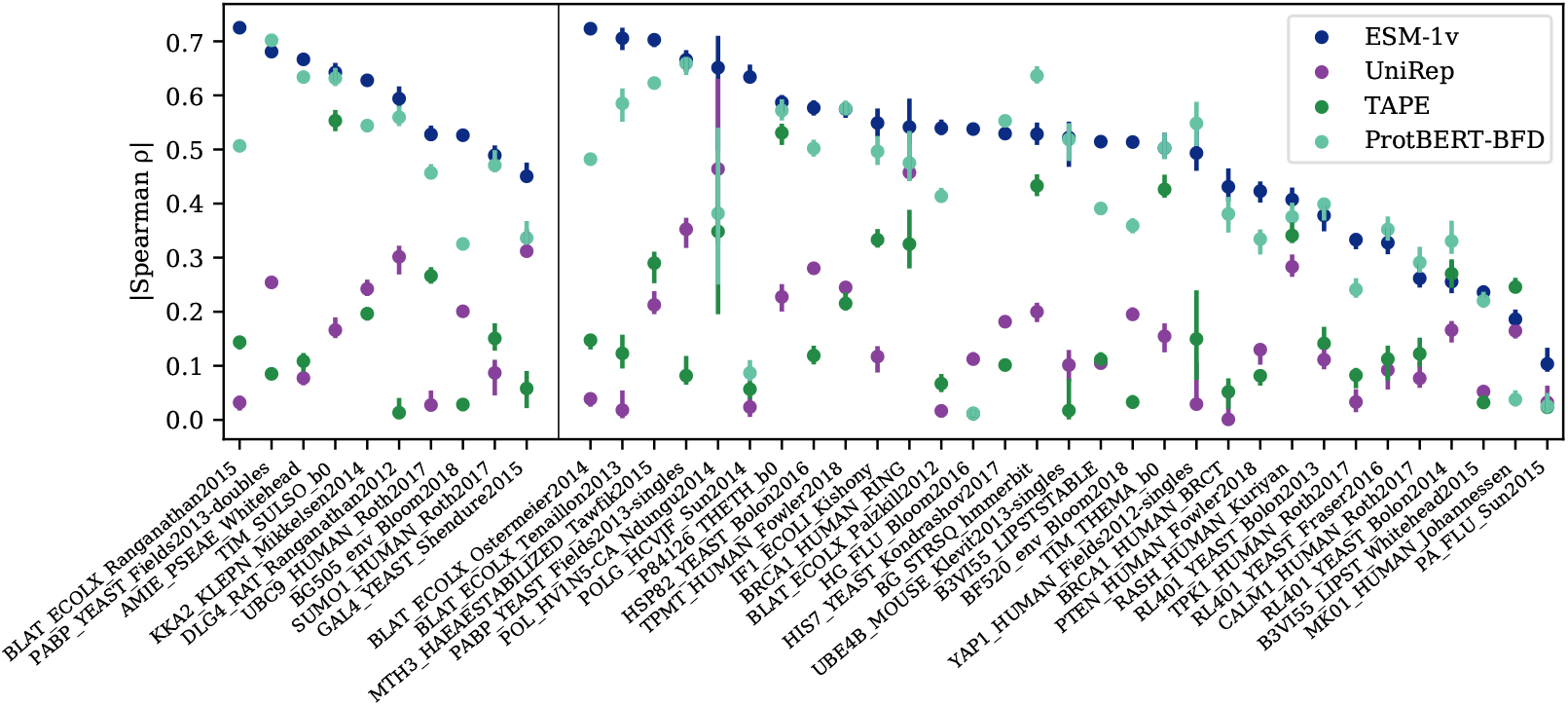
Zero-shot performance of ESM-1v compared to earlier protein language models on all 41 deep mutational scans. Points are |Spearman *ρ*| on each dataset, error bars show standard deviation of 20 bootstrapped samples. Validation proteins are shown to the left of the dividing line and test proteins to the right. ESM-1v is the best performing method on 30 of the 41 deep mutational scans.

This formulation assumes an additive model, consistent with the training objective. We show that this assumption is justified empirically by evaluating the model with different choices at the non-mutated positions. First, the wildtype sequence (Strategy b):

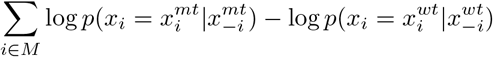

and the mutant sequence (Strategy c):

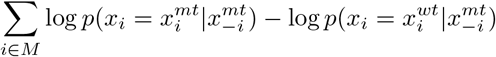

**Table 4:**
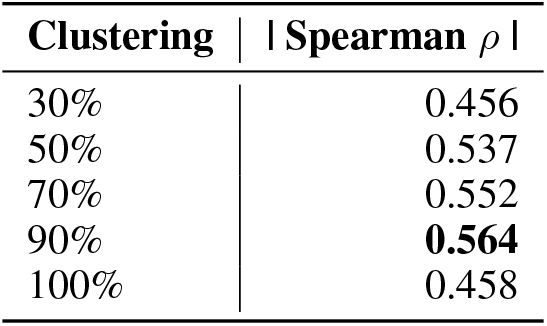
Average |Spearman *ρ*| on the single-mutation validation set after training a 650M parameter Transformer model for 170,000 updates on various sequence identity clusterings of Uniref.

| Clustering | \| Spearman $\rho$ \| |
| --- | --- |
| 30% | 0.456 |
| 50% | 0.537 |
| 70% | 0.552 |
| 90% | **0.564** |
| 100% | 0.458 |

**Table 5:** Benchmarking scoring schemes on the single-mutation validation set. The means across the validation set are listed. The masked marginal scheme performs best.

| Method | \| Spearman $\rho$ \| |
| --- | --- |
| Masked marginal | **0.582** |
| Mutant marginal | 0.578 |
| Wildtype marginal | 0.572 |
| Pseudo-likelihood | 0.552 |

Strategy (a), where we mask all positions at the same time, performs best on the PABP Yeast Doubles validation dataset (Table 7).

## Mutant marginal probability

This method is analogous to the wildtype marginal probability, except we use the mutant sequence instead.

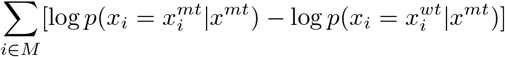

This method requires a single forward pass for every mutation.

## Wildtype marginal probability (1 forward pass)

In the fastest scheme, we perform a single forward pass using the wildtype sequence as input. For a set of mutations at positions *M*, the score is:

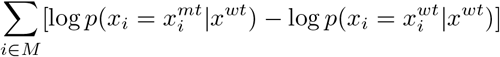

We find that the method performs well with a minor 1% decrease in absolute performance, while requiring very limited computational resources. The strong performance indicates that the masked language modeling objective causes the model to capture the fitness landscape of the protein in its outputs.

## Pseudolikelihood

Psuedolikelihood has been proposed in the literature as a method to score sequences using masked language models [86]. We compute the score as follows:

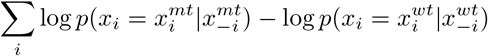

As mutation prediction is a ranking task and as the contribution from the second term is constant throughout the deep mutation scan (i.e. the wildtype sequence is always the same), we can safely drop it from the computation.

## A.1 Evaluation

We compare the methods described above on the validation set, finding that the masked marginal scheme performs best. To determine the specific mode of inference when multiple mutations are present, we examine each method on the “doubles” component of the PABP Yeast dataset finding the masked marginal (a) strategy performs best. This scoring method is used across the results.

**Table 6:** ESM-1v performs better when including the full protein sequence as listed in Uniprot, compared to using the seed sequence of the MSA corresponding to the deep mutational scan. Results on single-mutation validation set. The means across the validation set are listed. We experiment with a number of strategies for inference: (i) the consensus columns only; (ii) the aligned part of the query sequence; and (iii) the complete Uniprot sequence. The complete Uniprot sequence performs best, possibly because the model was pre-trained on complete Uniprot sequences. We use the MSA seed sequence from the MSAs released by [20] corresponding to the deep mutational scans.

| Input | Consensus columns only | \| Spearman $\rho$ \| |
| --- | --- | --- |
| MSA seed | Yes | 0.573 |
| MSA seed | No | 0.567 |
| Uniprot | N/A | **0.582** |

**Table 7:** Ablating scoring schemes on the PABP Yeast Doubles dataset. The masked marginal scheme performs best when masking all mutated sites together. Mean absolute Spearman *ρ* across the single-mutation validation tasks is reported.

| Method | \| Spearman $\rho$ \| |
| --- | --- |
| Masked marginal (a) | **0.692** |
| Masked marginal (b) | 0.482 |
| Masked marginal (c) | 0.483 |
| Mutant marginal | 0.694 |
| Wildtype marginal | 0.672 |
| Pseudo-likelihood | 0.608 |

## A.2 Evaluating ESM-1v on subsequences

DeepSequence, EVMutation, and the MSA Transformer use the consensus columns of a MSA as input. We construct MSAs using the seed sequences from the DeepSequence paper, which usually correspond to a subsequence of the protein capturing the domain where the deep mutational scan was performed.

Table 6 explores using the MSA seed sequence vs. the full Uniprot sequence for inference on the validation set. We find that the full Uniprot sequence performs best, possibly because the model was pre-trained on Uniprot sequences. We note in Figure Fig. 6 that the model captures some bias in the Uniprot dataset, for example that most proteins begin with a methionine (corresponding to the start codon).

## B Unsupervised fine-tuning ESM-1v

## Experimental setup

We assess a number of approaches for fine-tuning ESM-1v on task-specific MSAs. We evaluate modeling decisions by fine-tuning on tasks from the validation set and examining the mean change in Spearman *ρ* over the course of training. For efficiency, we compute Spearman *ρ* using the wildtype marginal strategy, as this requires just a single forward pass. After the final modeling decisions are selected, we train all models for 7500 updates and evaluate on all proteins using the masked marginals strategy. All models in this section were trained with a constant learning rate of 10^−5^ using the masked language modeling objective. For reference, the ESM-1v pre-training was performed with a target batch size of 1M tokens.

## Unsupervised fine-tuning baselines

The concept of unsupervised fine-tuning of an MSA has been previously proposed [21, 16]. Fig. 12 studies a basic fine-tuning setup on the consensus columns of the MSA. Each model is fine-tuned on a single MSA with a target batch size of 8192 tokens. We first observe that that models overfit quickly if the entire model is trained. This results in a decrease in Spearman *ρ* compared to initialization. As the fine-tuning is performed on the consensus columns of the MSA, we sought to regularize the model by fine-tuning only the embeddings. As a PSSM already captures information relevant to the task, we hypothesize that tuning the embeddings could capture similar information and boost performance. Similarly, we experiment with tuning only the layer normalizations, as these have also been recently shown to enable transfer to new tasks. In both cases, we found no improvement to the average Spearman *ρ*. We also assessed label smoothing and replacing the gap token with a mask token or a pad token finding no significant impact; for simplicity, we omit label smoothing and use the mask token for future experiments.

**Figure 9:**
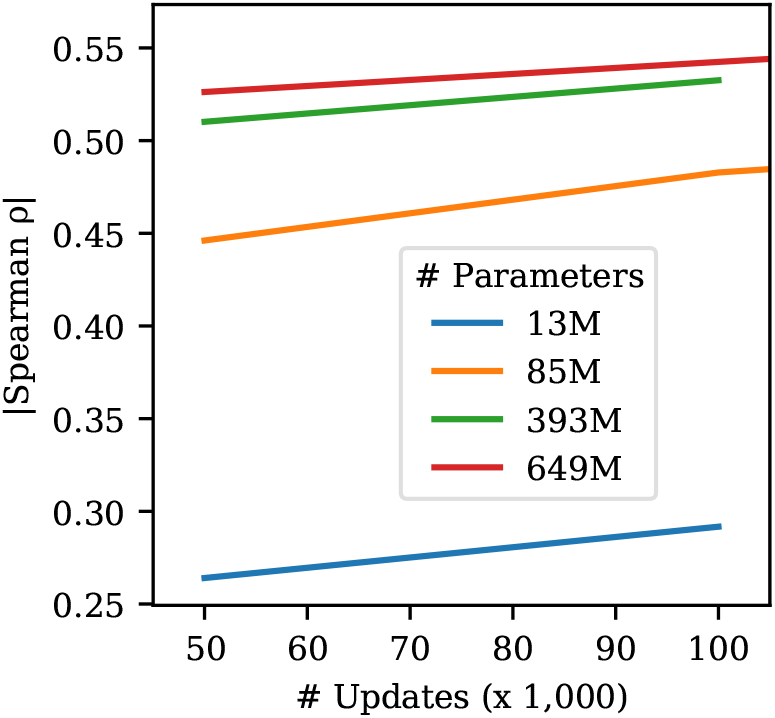
Larger models perform better on variant prediction. We trained four models of various scales, following the hyperparameters listed in Henighan et al. [27]. Results on single-mutation validation set.

**Table 8:** Subsampling strategies for MSA Transformer evaluated on the single-mutation validation set. Sequence reweighting performs best. When sampling methods are stochastic, 5 seeds are run and the mean and standard deviation is reported. With HHFilter, we run with the `-diff M` parameter and randomly subsample the output if more than `M` sequences are returned. We use a coverage parameter of 75 and a sequence identity parameter of 99. Mean absolute Spearman *ρ* across the single-mutation validation tasks is reported.

| MSA Subsample Strategy | Context size | \|Spearman $\rho$\| |
| --- | --- | --- |
| Diversity minimizing | 256 sequences | 0.255 |
| Random | 256 sequences | 0.535 ± 0.024 |
| HHFilter | 256 sequences | 0.550 ± 0.015 |
| Sequence reweighting | 256 sequences | 0.578 ± 0.005 |

## Minimal models

We also examine a set of minimal models, in which we freeze all parameters in the Transformer and learn a projection from the ESM-1v outputs onto a PSSM, taking the sum of the projection and the PSSM. We experiment with freezing the PSSM or allowing it to train. We did not see a change in Spearman *ρ* of more than 0.01.

## Spiked unsupervised fine-tuning

Next, we examine a new strategy, which we call `spiked fine-tuning`. In spiked fine-tuning, we regularize the fine-tuning by continuing to spike pretraining sequences into the fine-tuning batch. In this setting, we train on the entire MSA, including non-consensus positions. We find that spiked fine-tuning with a small ratio (0.01) of MSA tokens to pre-training tokens performs best and enables training of all parameters without overfitting.

The final models were trained for 7500 updates using spiked fine-tuning with a batch size of 500k tokens. To produce an ensemble, we perform the fine-tuning scheme on five models that were pre-trained with different seeds. Each model was also fine-tuned with a unique seed.

**Figure 10:**
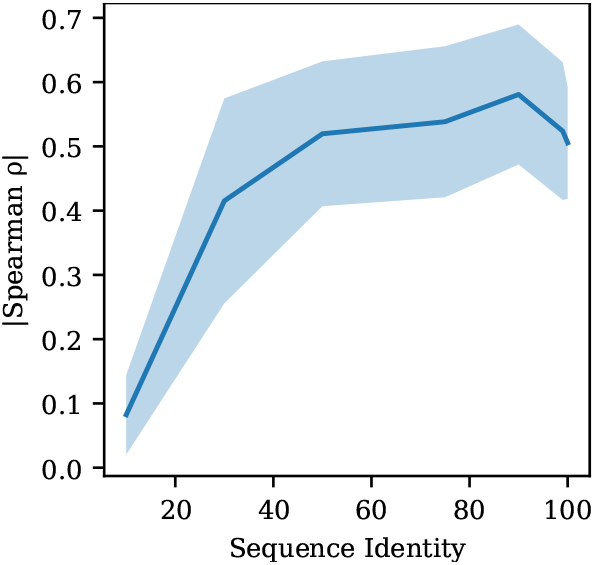
Filtering sequences with high sequence identity to the query improves performance. The curve illustrates mean ± standard deviation across the 9 validation proteins. HHFilter is used to filter the MSAs with coverage of 75 and various sequence identity values as shown on x-axis. After filtering, 384 sequences are sampled for inference. Each sequence identity value *s* refers to using sequences with no more than *s*% sequence identity to the seed sequence. The MSA Transformer appears to primarily use sequences that are close to the seed sequence, yet performance drops if sequences that are *too similar* remain in the MSA. Results are broken down across the single-mutation validation set in Table 9.

**Table 9:** Filtering MSAs from the single-mutation validation set with HHFilter coverage 75 and various sequence identity values. Filtering sequences to an identity threshold of 75% or 90% consistently performs best. The Spearman rank correlation between MSA Transformer predictions and experimental data is shown for each deep mutational scan.

| Sequence Identity (%) | 10 | 30 | 50 | 75 | 90 | 99 | 100 |
| --- | --- | --- | --- | --- | --- | --- | --- |
| AMIE_PSEAE_Whitehead | 0.025 | 0.461 | 0.467 | 0.365 | **0.665** | 0.654 | 0.622 |
| BG505_env_Bloom2018 | 0.055 | 0.055 | 0.417 | **0.482** | 0.452 | 0.450 | 0.457 |
| BLAT_ECOLX_Ranganathan2015 | 0.060 | 0.630 | 0.745 | 0.776 | **0.795** | 0.662 | 0.478 |
| DLG4_RAT_Ranganathan2012 | 0.008 | 0.431 | 0.416 | 0.431 | **0.457** | 0.418 | 0.400 |
| GAL4_YEAST_Shendure2015 | 0.080 | 0.287 | 0.366 | 0.441 | **0.576** | 0.542 | 0.388 |
| SUMO1_HUMAN_Roth2017 | 0.131 | 0.430 | 0.516 | **0.541** | 0.500 | 0.492 | 0.495 |
| TIM_SULSO_b0 | 0.026 | 0.581 | 0.633 | 0.625 | **0.649** | 0.324 | 0.632 |
| UBC9_HUMAN_Roth2017 | 0.165 | 0.376 | 0.557 | **0.583** | 0.494 | 0.555 | 0.475 |
| KKA2_KLEPN_Mikkelsen2014 | 0.192 | 0.484 | 0.560 | 0.601 | **0.637** | 0.616 | 0.602 |

## C Datasets

## C.1 Evaluation Tasks

We evaluate models on a set of 41 deep mutational scans collected by Riesselman et al. [20], which comprise a variety of tasks assessing a diverse set of proteins. Across tasks, the experimental data differ widely in the functions tested and in the experimental measurements performed. Of the 41 datasets, 37 are single-mutation only, 1 is double-mutation only, and the rest contain a variable number of mutations per sequence between 1 and 28. The median number of mutations is 2979, and the average is 16822; the smallest dataset has 37 mutations, and the largest has 496137. We randomly select 9 single-mutation experiments as a validation set. We also ablate the multiple mutation scoring approach on the double mutations from the PABP Yeast deep mutational scan. We exclude the 10 tasks used for validation and ablations from the test set. These datasets are reported in results for the full set. While the original compilation has 43 datasets, we exclude the tRNA (which is not a protein) and the toxin-antitoxin complex (which comprises multiple proteins).

We treat each deep mutational scanning dataset as a separate prediction task, scoring each of the variants in the dataset with the model. We evaluate performance by comparing the scores with the experimental measurements using Spearman rank correlation. Results are broken out between the test set, which excludes the validation set, as well as the full set of 41 datasets. All ablations are performed on the single mutant validation set or the PABP Yeast doubles experiment. Only the final models are evaluated on the test set.

**Figure 11:**
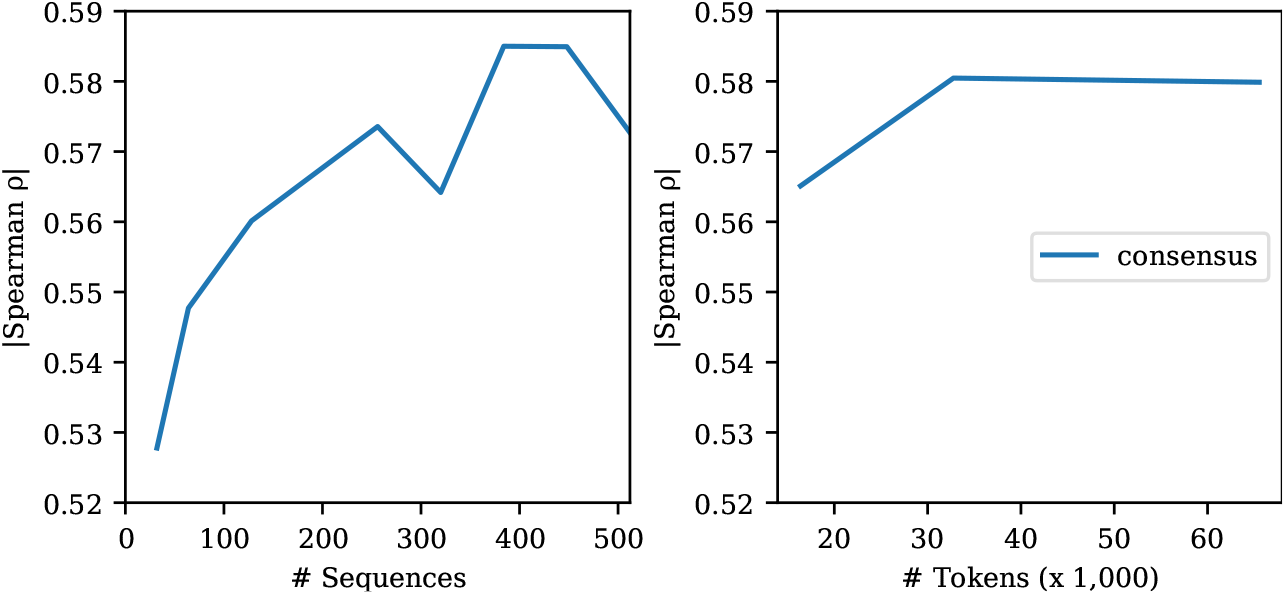
Few-shot performance of the MSA Transformer is robust to the number of sequences used for inference. **Left**: Varying the number of sequences used in inference. **Right**: Varying the number of tokens used for inference. Since the number of sequences in each MSA varies, we assess the effect of fixing the total number of tokens sampled from each MSA and drawing the corresponding number of sequences to fill the context. Results on single-mutation validation set.

**Figure 12:**
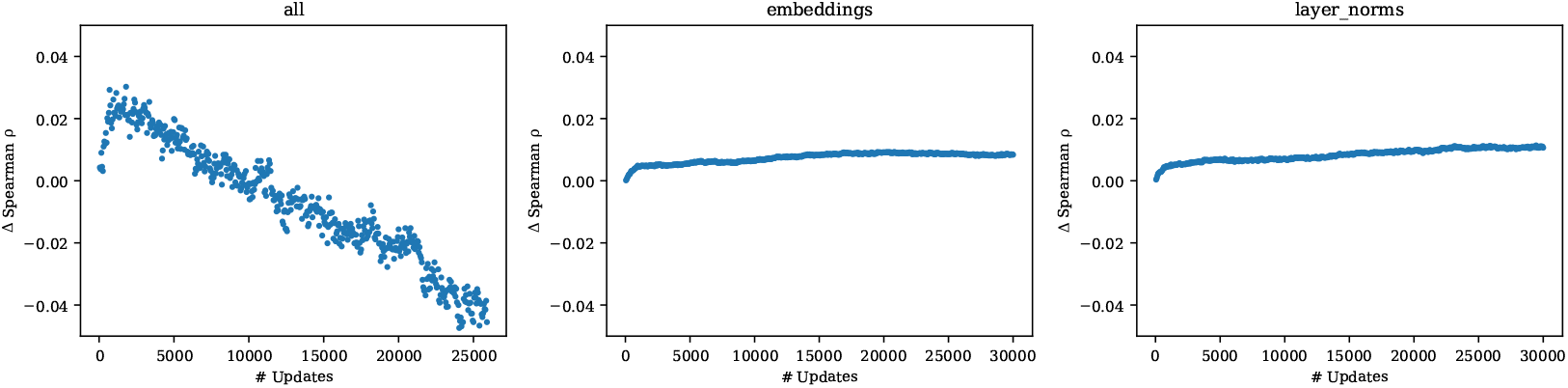
Unsupervised fine-tuning baselines. Mean change in Spearman *ρ* across 9 models trained on the single-mutation validation set tasks. The title of each plot denotes the parameters that are trained. We find that fine-tuning the entire model results in overfitting, but limiting the training to just the embeddings or just the layer norms does not improve performance with respect to the pre-trained initialization. The choice of gap token and label smoothing has limited effect.

## C.2 Pre-training datasets

For the clustering sweep in Fig. 4, we use the Uniref50 and Uniref90 databases from the `2020_03` release of Uniref [24], a publicly available database of proteins, clustered respectively to 50% and 90% sequence identity. For the 30% sequence identity dataset, we use Uniclust30 `2020_03` [89]. For the 70% sequence identity dataset, we Uniref100 is hierarchically clustered to the 90%, then 70% sequence identity level. MMseqs settings are those used by Uniref: 80% overlap with longest sequnece in the cluster, which translates to `mmseqs-cluster -min-seq-id 90,70 -cov-mode 0 -alignment-mode 3 -c 0.8`. In order to compute pre-training perplexities on a heldout validation set, we randomly select 1% of sequences each from Uniref30, Uniref50, and Uniref90. We then exclude sequences that are similar to the validation sequences by removing all sequences found with MMSeqs search (`-min-seq-id 0.xx`) for validation set xx. We use the most sensitive settings in MMSeqs `-alignment-mode 3 -max-seqs 300 -s 7`, taking the train set as the query database and the validation set as the target database. We use settings `-c 0.8 -cov-mode 0` to match the settings of Uniref. Pretraining perplexities on the validation sets are reported in Table 10.

**Figure 13:**
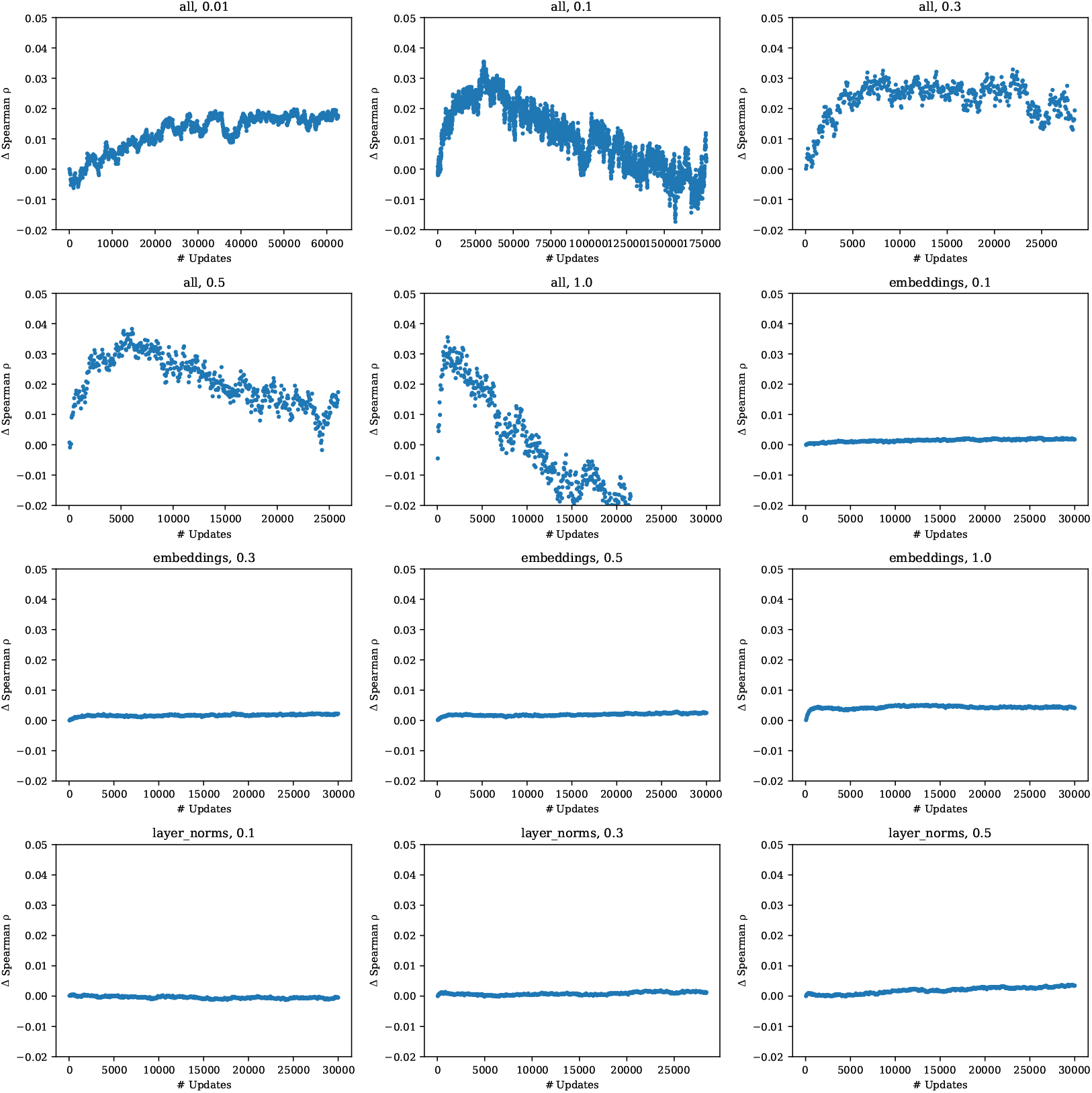
Spiked unsupervised fine-tuning. Mean change in Spearman *ρ* across 9 models trained on the single-mutation validation tasks. The title of each plot denotes the parameters that are trained; and the ratio of MSA tokens to pre-training tokens. We find that a small ratio performs well and reduces the tendency for the model to overfit, while preserving strong performance. Performance is not improved if the fine-tuning is limited to just the embeddings or just the layer norms.

**Figure 14:**
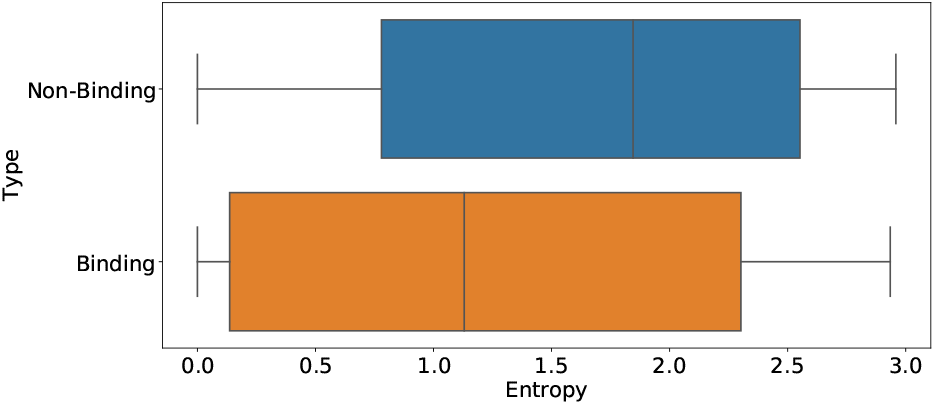
Box plot comparing entropy scores for binding vs non-binding positions in structures labeled in the Provis validation dataset (as described in Appendix B.4 of [36]). A Welch’s *t*-test determines that the difference between the two means is statistically significant (p < 0.01).

**Figure 15:**
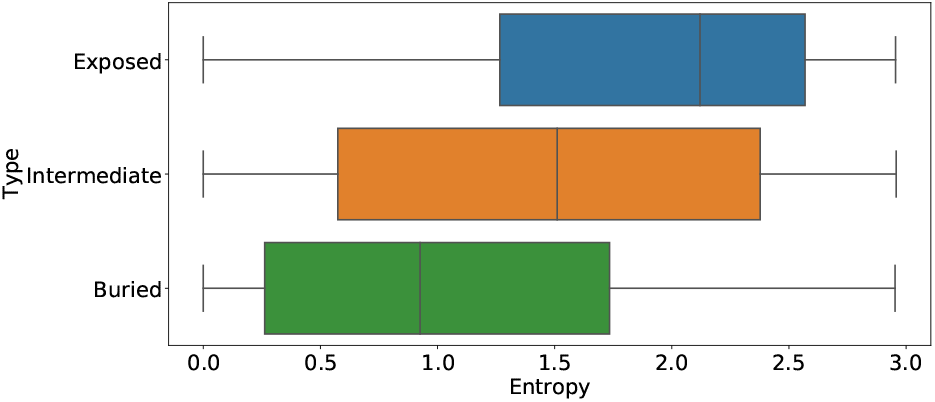
Box plot comparing entropy scores across residue depths in structures from the trRosetta dataset. Residue depths are categorized based on the number of neighboring residues with C-beta distance<10 angstroms. (exposed≤16, buried≥24 [88]). A one way Anova test determines that the differences between all three means are statistically significant (p < 0.01).

## C.3 Baselines

The MSAs used for training DeepSequence and EVMutation are generated from the 2017-10 version of Uniref100, whereas the models we study are trained on sequences from Uniref90 2020-03. In the case of MSA Transformer, the model is pre-trained on the 2018-03 Uniref, but we use 2020-03 MSAs for inference. In order to provide a fair comparison, we regenerate MSAs against the 2020-03 Uniref according to the methodology in Hopf et al. [4] and retrain EVMutation (replication) and DeepSequence (replication) on these datasets using their open-source codebases. For the viral proteins `BF520_env_Bloom2018, BG505_env_Bloom2018, HG_FLU_Bloom2016, PA_FLU_Sun2015, POLG_HCVJF_Sun2014, POL_HV1N5-CA_Ndungu2014`, we compute the sequence weights with *θ* = 0.01 (versus default *θ* = 0.2) following Riesselman et al. [20]. In the replication of the DeepSequence ensemble, for `BF520_env_Bloom2018, BG505_env_Bloom2018`, one of the five runs failed so we reran with a different random seed.

## C.4 Validation and test set

The single-mutation validation set consists of the following deep mutational scans: AMIE_PSEAE_Whitehead, BG505_env_Bloom2018, BLAT_ECOLX_Ranganathan2015, BRCA1_HUMAN_RING, DLG4_RAT_Ranganathan2012, GAL4_YEAST_Shendure2015, POLG_HCVJF_Sun2014, SUMO1_HUMAN_Roth2017, TIM_SULSO_b0, UBC9_HUMAN_Roth2017, KKA2_KLEPN_Mikkelsen2014.

For ablations studies with multiple mutations the following dataset is used: `PABP_YEAST_Fields2013-doubles`

The test set consists of the following deep mutational scans: B3VI55_LIPSTSTABLE, B3VI55_LIPST_Whitehead2015, BF520_env_Bloom2018, BG_STRSQ_hmmerbit, BLAT_ECOLX_Ostermeier2014, BLAT_ECOLX_Palzkill2012, BLAT_ECOLX_Tenaillon2013, BRCA1_HUMAN_BRCT, CALM1_HUMAN_Roth2017, HG_FLU_Bloom2016, HIS7_YEAST_Kondrashov2017, HSP82_YEAST_Bolon2016, IF1_ECOLI_Kishony, MK01_HUMAN_Johannessen, MTH3_HAEAESTABILIZED_Tawfik2015, P84126_THETH_b0, PABP_YEAST_Fields2013-singles, PA_FLU_Sun2015, POL_HV1N5-CA_Ndungu2014, PTEN_HUMAN_Fowler2018, RASH_HUMAN_Kuriyan, RL401_YEAST_Bolon2013, RL401_YEAST_Bolon2014, RL401_YEAST_Fraser2016, TIM_THEMA_b0, TPK1_HUMAN_Roth2017, TPMT_HUMAN_Fowler2018, UBE4B_MOUSE_Klevit2013-singles, YAP1_HUMAN_Fields2012-singles.

**Figure 16:**
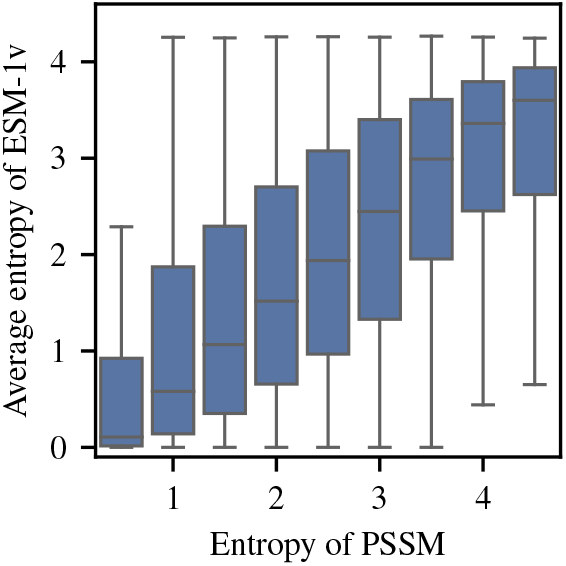
Entropy of PSSM versus ESM-1v predicted entropy on the trRosetta dataset. PSSM entropy determines the level of conservation at a given position in a protein family. ESM-1v entropy is well correlated with PSSM entropy (Pearson’s *r* = 0.44), suggesting the model is able to identify conserved positions.

**Figure 17:**
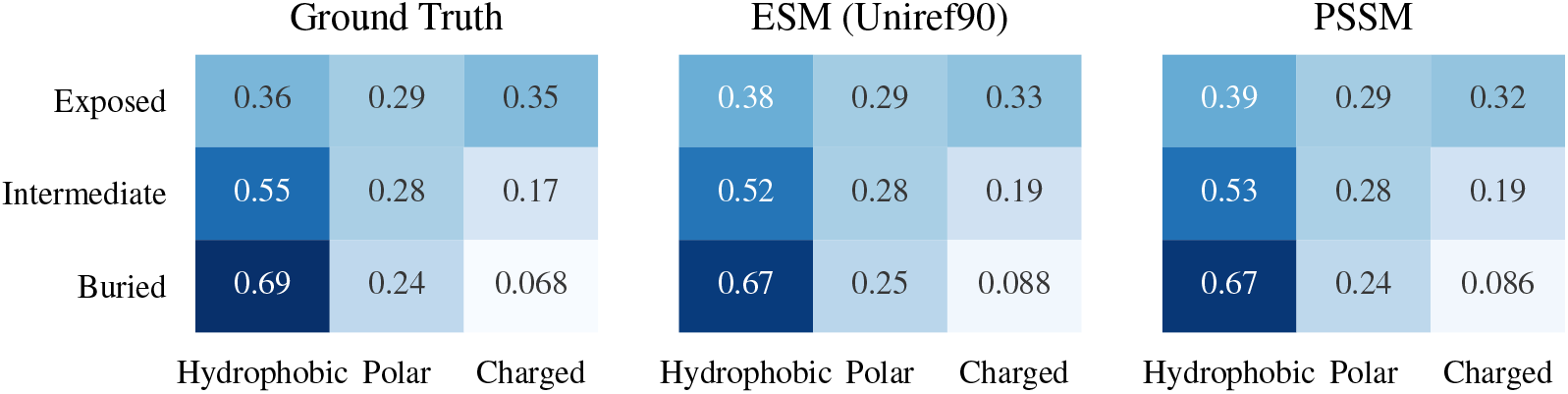
Predicted distribution of hydrophobic, polar and charged amino acids at the surface and core of proteins in the trRosetta dataset. We compare to the actual proportion in the protein structure. We classify residues into buried, intermediate or exposed by residue depths based on the number of neighboring residues with C-beta distance < 10 angstroms (exposed ≤ 16, buried ≥ 24) [88]. ESM-1v and PSSM both see increased hydrophobicity predictions for buried residues, in correspondence with the ground truth data. Predicted probabilities are produced by introducing a mask token at each position.

## D Methodology

## D.1 Model selection

ESM-1b and MSA Transformer model checkpoints are selected based on performance on the single mutation validation set. Open sourced checkpoints are used for ESM-1b and other protein language model baselines.

## D.2 Treatment of synonymous mutations

Synonymous mutations are mutations in DNA that do not change the protein sequence that is expressed. The deep mutational scanning datasets that we evaluate here can therefore include DNA mutations that do not change the protein sequence itself. Synonymous mutations are excluded from results.

## D.3 Bootstraps

To compute bootstraps for the pointplots, we randomly resample each deep mutational scan (with replacement) and compute the Spearman *ρ* between the experimental data and model predictions.

## D.4 Average calibration error

The standard expected calibration error (ECE) performs poorly for highly imbalanced data [90]. Following Neumann et al. [90] and Nixon et al. [91] we adapt average calibration error for the multi-class setting as follows:

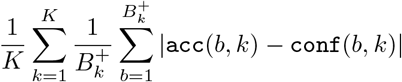

where *K* is the number of classes, 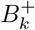 is the number of non-empty bins for class *k*, and `acc` and `conf` are the accuracy and confidence for bin *b* and class *k*.

## E Performance by MSA depth

We examine the relationship between the number of related sequences in the pre-training set and performance on the task. We use Jackhmmer [92] version 3.3.1 with a bitscore threshold of 27 and 8 iterations to construct MSAs from the ESM-1v training set. We do not observe a strong correlation between MSA depth and the observed absolute value of Spearman *ρ* (Figure Fig. 19).

## F Compute costs

ESM-1v models are pre-trained for 6 days on 64 V100 GPUs. Weights for the MSA Transformer were retrieved from the open-source repository released by the authors; the model was pre-trained for 13 days on 128 V100 GPUs. Once trained, the models can be used directly for function prediction tasks. Forward inference is efficient, meaning that for applications of the models, the additional compute is minimal. In total, five ESM-1v models were trained on various Uniref clustering thresholds to five different levels: 30%, 50%, 70%, 90%, and 100%. For the 90% sequence identity level, five total models with different random seeds were trained, for use in an ensemble. As illustrated in Fig. 7, inference is inexpensive by comparison. Batch inference was performed with preemptible, short (shorter than one hour), single V100 GPU jobs on a shared compute cluster.

**Table 10:** Perplexities on heldout pre-training validation sequences after training a 650M parameter Transformer model for 170,000 updates on various sequence identity clusterings of Uniref.

| Clustering | Valid (30%) | Valid (50%) | Valid (90%) |
| --- | --- | --- | --- |
| 30% | 8.93 | 8.33 | 7.29 |
| 50% | 8.90 | 7.77 | 6.27 |
| 70% | 9.05 | 7.80 | 5.85 |
| 90% | 9.37 | 8.10 | 5.56 |
| 100% | 9.89 | 8.65 | 6.05 |

**Figure 18:**
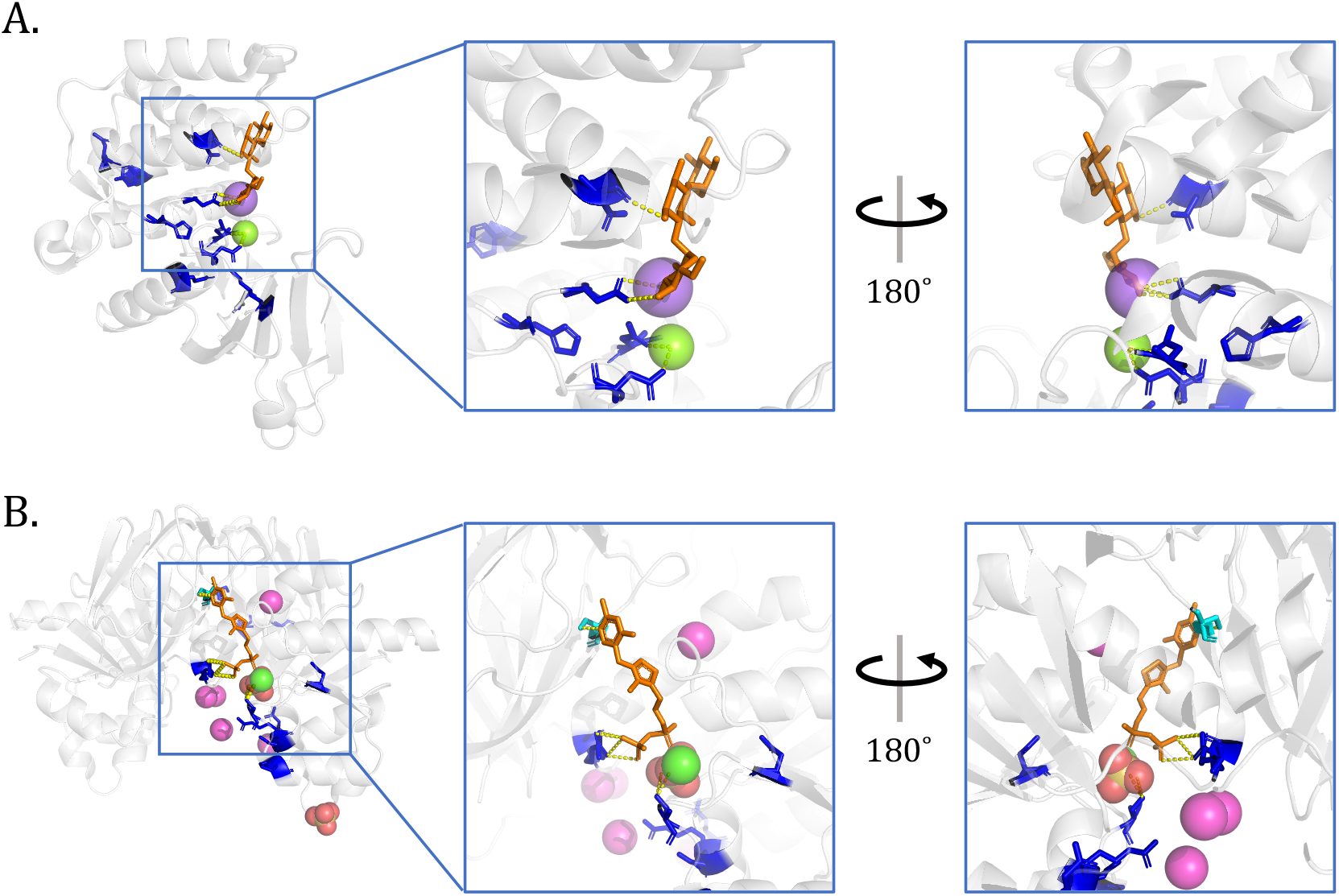
ESM-1v accurately captures functional properties. Further examples. The ten positions with lowest predicted entropy highlighted in blue. **(A)** Kanamycin kinase APH(3’)-II (pdbid: 1ND4 [93]). The highlighted residues interact with the kanamycin aminoglycoside, as well as the magnesium and sodium ions. **(B)** Thiamin pyrophosphokinase 1 (pdbid: 3S4Y). Residue 216 is one of the 10 lowest entropy residues, and we highlight it on the other chain (in cyan) to show both chains of the dimer interacting with the thiamine diphosphate.

**Figure 19:**
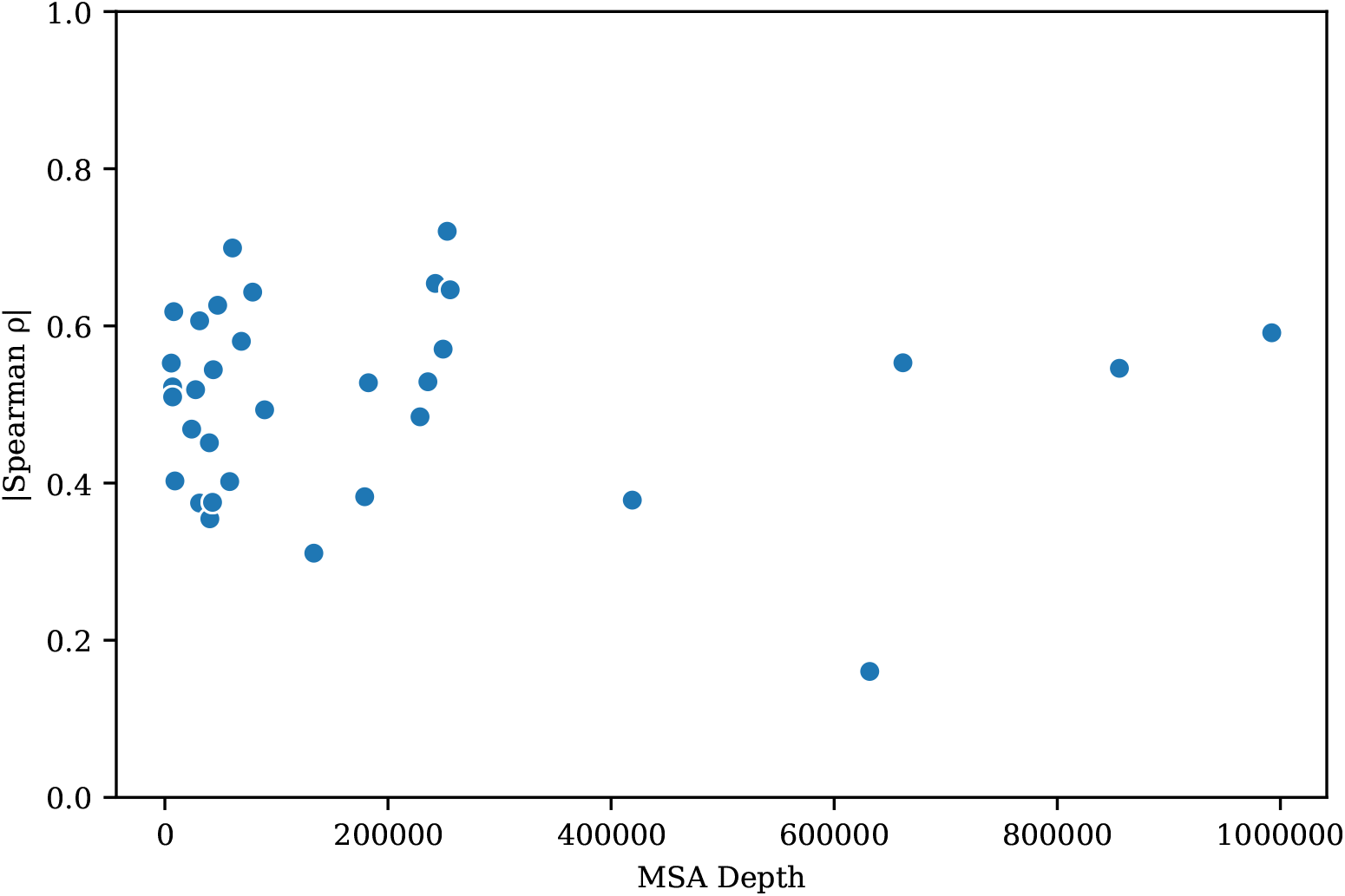
Relation between MSA depth and zero-shot performance of ESM-1v. We use JackHMMer [92] version 3.3.1 with a bitscore threshold of 27 and 8 iterations to construct MSAs from the ESM-1v training set. We do not observe a strong correlation between MSA depth and the observed |Spearman *ρ*|.

## Notes

https://github.com/facebookresearch/esm

### Competing Interest Statement

The authors have declared no competing interest.

## References

[1] Charles J Epstein, Robert F Goldberger, and Christian B Anfinsen. The genetic control of tertiary protein structure: studies with model systems. In Cold Spring Harbor symposia on quantitative biology, volume 28, pages 439–449. Cold Spring Harbor Laboratory Press, 1963.

[2] Douglas M Fowler and Stanley Fields. Deep mutational scanning: a new style of protein science. Nature methods, 11(8):801, 2014.

[3] Matteo Figliuzzi, Hervé Jacquier, Alexander Schug, Oliver Tenaillon, and Martin Weigt. Co-evolutionary landscape inference and the context-dependence of mutations in beta-lactamase tem-1. Molecular biology and evolution, 33(1):268–280, 2016.

[4] Thomas A. Hopf, John B. Ingraham, Frank J. Poelwijk, Charlotta P.I. Schärfe, Michael Springer, Chris Sander, and Debora S. Marks. Mutation effects predicted from sequence co-variation. Nature Biotechnology, 35(2):128–135, feb 2017. ISSN 15461696. doi: 10.1038/nbt.3769.

[5] C Yanofsky, V Horn, and D Thorpe. Protein Structure Relationships Revealed By Mutational Analysis. Science (New York, N.Y.), 146(3651):1593–4, 12 1964. ISSN 0036-8075. URL http://www.ncbi.nlm.nih.gov/pubmed/14224506.

[6] D. Altschuh, A.M. Lesk, A.C. Bloomer, and A. Klug. Correlation of co-ordinated amino acid substitutions with function in viruses related to tobacco mosaic virus. Journal of Molecular Biology, 193(4):693–707, 1987. ISSN 0022-2836. doi: 10.1016/0022-2836(87)90352-4.

[7] D Altschuh, T Vernet, P Berti, D Moras, and K Nagai. Coordinated amino acid changes in homologous protein families. Protein Engineering, Design and Selection, 2(3):193–199, 1988.

[8] Ulrike Göbel, Chris Sander, Reinhard Schneider, and Alfonso Valencia. Correlated mutations and residue contacts in proteins. Proteins: Structure, Function, and Genetics, 18(4):309–317, 4 1994. ISSN 0887-3585. doi: 10.1002/prot.340180402. URL http://www.ncbi.nlm.nih.gov/pubmed/8208723http://doi.wiley.com/10.1002/prot.340180402.

[9] Alec Radford, Jeff Wu, Rewon Child, David Luan, Dario Amodei, and Ilya Sutskever. Language models are unsupervised multitask learners. 2019.

[10] Tom B. Brown, Benjamin Mann, Nick Ryder, Melanie Subbiah, Jared Kaplan, Prafulla Dhariwal, Arvind Neelakantan, Pranav Shyam, Girish Sastry, Amanda Askell, Sandhini Agarwal, Ariel Herbert-Voss, Gretchen Krueger, Tom Henighan, Rewon Child, Aditya Ramesh, Daniel M. Ziegler, Jeffrey Wu, Clemens Winter, Christopher Hesse, Mark Chen, Eric Sigler, Mateusz Litwin, Scott Gray, Benjamin Chess, Jack Clark, Christopher Berner, Sam McCandlish, Alec Radford, Ilya Sutskever, and Dario Amodei. Language models are few-shot learners. CoRR, abs/2005.14165, 2020. URL https://arxiv.org/abs/2005.14165.

[11] Alec Radford, Jong Wook Kim, Chris Hallacy, Aditya Ramesh, Gabriel Goh, Sandhini Agarwal, Girish Sastry, Amanda Askell, Pamela Mishkin, Jack Clark, Gretchen Krueger, and Ilya Sutskever. Learning transferable visual models from natural language supervision, 2021.

[12] Alexander Rives, Joshua Meier, Tom Sercu, Siddharth Goyal, Zeming Lin, Jason Liu, Demi Guo, Myle Ott, C. Lawrence Zitnick, Jerry Ma, and Rob Fergus. Biological Structure and Function Emerge from Scaling Unsupervised Learning to 250 Million Protein Sequences. bioRxiv, page 622803, 4 2020. doi: 10.1101/622803. URL https://www.biorxiv.org/content/10.1101/622803v3.

[13] Roshan Rao, Jason Liu, Robert Verkuil, Joshua Meier, John F. Canny, Pieter Abbeel, Tom Sercu, and Alexander Rives. MSA Transformer. bioRxiv, page 2021.02.12.430858, feb 2021. doi: 10.1101/2021.02.12.430858.

[14] Ahmed Elnaggar, Michael Heinzinger, Christian Dallago, Ghalia Rihawi, Yu Wang, Llion Jones, Tom Gibbs, Tamas Feher, Christoph Angerer, Martin Steinegger, Debsindhu Bhowmik, and Burkhard Rost. ProtTrans: Towards Cracking the Language of Life’s Code Through Self-Supervised Deep Learning and High Performance Computing. bioRxiv, 7 2020. URL http://arxiv.org/abs/2007.06225.

[15] Chloe Hsu, Hunter Nisonoff, Clara Fannjiang, and Jennifer Listgarten. Combining evolutionary and assay-labelled data for protein fitness prediction. bioRxiv, page 2021.03.28.437402, mar 2021. doi: 10.1101/2021.03.28.437402.

[16] Surojit Biswas, Grigory Khimulya, Ethan C Alley, Kevin M Esvelt, and George M Church. Low-N protein engineering with data-efficient deep learning. bioRxiv, page 2020.01.23.917682, 2020. doi: 10.1101/2020.01.23.917682.

[17] Christoph H Lampert, Hannes Nickisch, and Stefan Harmeling. Learning to detect unseen object classes by between-class attribute transfer. In 2009 IEEE Conference on Computer Vision and Pattern Recognition, pages 951–958. IEEE, 2009.

[18] Hugo Larochelle, Dumitru Erhan, and Yoshua Bengio. Zero-data learning of new tasks. In AAAI, volume 1, page 3, 2008.

[19] Ramesh A, Pavlov M, Goh G, Gray S, Voss C, Radford A, Chen M, and Sutskever I. Zero-shot text-to-image generation.

[20] Adam J. Riesselman, John B. Ingraham, and Debora S. Marks. Deep generative models of genetic variation capture the effects of mutations. Nature Methods, 15(10):816–822, 10 2018. ISSN 15487105. doi: 10.1038/s41592-018-0138-4.

[21] Ethan C. Alley, Grigory Khimulya, Surojit Biswas, Mohammed AlQuraishi, and George M. Church. Unified rational protein engineering with sequence-only deep representation learning. Nature Methods, 12:1315–1322, 3 2019. ISSN 15487105. doi: 10.1101/589333. URL https://www.biorxiv.org/content/10.1101/589333v1.

[22] Roshan Rao, Nicholas Bhattacharya, Neil Thomas, Yan Duan, Xi Chen, John Canny, Pieter Abbeel, and Yun S. Song. Evaluating Protein Transfer Learning with TAPE. In Neural Information Processing Systems. Cold Spring Harbor Laboratory, 6 2019. doi: 10.1101/676825. URL https://doi.org/10.1101/676825http://arxiv.org/abs/1906.08230.

[23] Robert D. Finn, Alex Bateman, Jody Clements, Penelope Coggill, Ruth Y. Eberhardt, Sean R. Eddy, Andreas Heger, Kirstie Hetherington, Liisa Holm, Jaina Mistry, Erik L.L. Sonnhammer, John Tate, and Marco Punta. Pfam: The protein families database, 1 2014. ISSN 03051048. URL https://www.ncbi.nlm.nih.gov/pmc/articles/PMC3965110/.

[24] Baris E. Suzek, Hongzhan Huang, Peter McGarvey, Raja Mazumder, and Cathy H. Wu. UniRef: Comprehensive and non-redundant UniProt reference clusters. Bioinformatics, 23(10):1282–1288, 5 2007. ISSN 13674803. doi: 10.1093/bioinformatics/btm098. URL http://www.uniprot.org.

[25] Jacob Devlin, Ming-Wei Chang, Kenton Lee, and Kristina Toutanova. BERT: Pre-training of Deep Bidirectional Transformers for Language Understanding. In Proceedings of the 2019 Conference of the North {A}merican Chapter of the Association for Computational Linguistics: Human Language Technologies, Volume 1 (Long and Short Papers), pages 4171–4186, Minneapolis, Minnesota, 6 2019. Association for Computational Linguistics. doi: 10.18653/v1/N19-1423. URL http://arxiv.org/abs/1810.04805.

[26] Roshan Rao, Joshua Meier, Tom Sercu, Sergey Ovchinnikov, and Alexander Rives. Transformer protein language models are unsupervised structure learners. ICLR, page 2020.12.15.422761, 12 2021. doi: 10.1101/2020.12.15.422761.

[27] Tom Henighan, Jared Kaplan, Mor Katz, Mark Chen, Christopher Hesse, Jacob Jackson, Heewoo Jun, Tom B. Brown, Prafulla Dhariwal, Scott Gray, Chris Hallacy, Benjamin Mann, Alec Radford, Aditya Ramesh, Nick Ryder, Daniel M. Ziegler, John Schulman, Dario Amodei, and Sam McCandlish. Scaling laws for autoregressive generative modeling. CoRR, abs/2010.14701, 2020. URL https://arxiv.org/abs/2010.14701.

[28] Faruck Morcos, Andrea Pagnani, Bryan Lunt, Arianna Bertolino, Debora S. Marks, Chris Sander, Riccardo Zecchina, José N. Onuchic, Terence Hwa, and Martin Weigt. Direct-coupling analysis of residue coevolution captures native contacts across many protein families. Proceedings of the National Academy of Sciences of the United States of America, 108(49):E1293–E1301, 12 2011. ISSN 00278424. doi: 10.1073/pnas.1111471108.

[29] KM Reinish, L Chen, GL Verdine, and WN Lipscomb. The crystal structure of haeiii methyltransferase covalently complexed to dna: an extrahelical cytosine and rearranged base pairing. Cell, 82(1):143–153, 1995.

[30] Michael Hennig, Beatrice Darimont, Reinhard Sterner, Kasper Kirschner, and Johan N Jansonius. 2.0 å structure of indole-3-glycerol phosphate synthase from the hyperthermophile sulfolobus solfataricus: possible determinants of protein stability. Structure, 3(12):1295–1306, 1995.

[31] Doris Forst, Wolfram Welte, Thomas Wacker, and Kay Diederichs. Structure of the sucrose-specific porin scry from salmonella typhimurium and its complex with sucrose. Nature structural biology, 5(1):37–46, 1998.

[32] Jianyi Yang, Ivan Anishchenko, Hahnbeom Park, Zhenling Peng, Sergey Ovchinnikov, David Baker, and John Harvard. Improved protein structure prediction using predicted inter-residue orientations. bioRxiv, page 846279, 2019. doi: 10.1101/846279. URL https://www.biorxiv.org/content/10.1101/846279v1.

[33] Tristan Bepler and Bonnie Berger. Learning protein sequence embeddings using information from structure, 2 2019. URL http://arxiv.org/abs/1902.08661https://arxiv.org/abs/1902.08661.

[34] Amy X Lu, Haoran Zhang, Marzyeh Ghassemi, and Alan Moses. Self-Supervised Contrastive Learning of Protein Representations By Mutual Information Maximization. bioRxiv, page 2020.09.04.283929, 9 2020. doi: 10.1101/2020.09.04.283929. URL https://doi.org/10.1101/2020.09.04.283929.

[35] Ali Madani, Bryan McCann, Nikhil Naik, Nitish Shirish Keskar, Namrata Anand, Raphael R. Eguchi, Po-Ssu Huang, and Richard Socher. ProGen: Language Modeling for Protein Generation. bioRxiv, 3 2020. URL http://arxiv.org/abs/2004.03497.

[36] Jesse Vig, Ali Madani, Lav R. Varshney, Caiming Xiong, Richard Socher, and Nazneen Fatema Rajani. BERTology Meets Biology: Interpreting Attention in Protein Language Models. bioRxiv, page 2020.06.26.174417, 6 2020. doi: 10.1101/2020.06.26.174417. URL http://arxiv.org/abs/2006.15222.

[37] Kevin K. Yang, Zachary Wu, and Frances H. Arnold. Machine-learning-guided directed evolution for protein engineering, aug 2019. ISSN 15487105.

[38] Vanessa E. Gray, Ronald J. Hause, Jens Luebeck, Jay Shendure, and Douglas M. Fowler. Quantitative Missense Variant Effect Prediction Using Large-Scale Mutagenesis Data. Cell Systems, 6(1):116–124.e3, jan 2018. ISSN 24054720. doi: 10.1016/j.cels.2017.11.003.

[39] Ivan A. Adzhubei, Steffen Schmidt, Leonid Peshkin, Vasily E. Ramensky, Anna Gerasimova, Peer Bork, Alexey S. Kondrashov, and Shamil R. Sunyaev. A method and server for predicting damaging missense mutations, apr 2010. ISSN 15487091.

[40] Nilah M Ioannidis, Joseph H Rothstein, Vikas Pejaver, Sumit Middha, Shannon K McDonnell, Saurabh Baheti, Anthony Musolf, Qing Li, Emily Holzinger, Danielle Karyadi, et al. Revel: an ensemble method for predicting the pathogenicity of rare missense variants. The American Journal of Human Genetics, 99(4):877–885, 2016.

[41] Kaitlin E Samocha, Jack A Kosmicki, Konrad J Karczewski, Anne H O’Donnell-Luria, Emma Pierce-Hoffman, Daniel G MacArthur, Benjamin M Neale, and Mark J Daly. Regional missense constraint improves variant deleteriousness prediction. BioRxiv, page 148353, 2017.

[42] Martin Kircher, Daniela M Witten, Preti Jain, Brian J O’Roak, Gregory M Cooper, and Jay Shendure. A general framework for estimating the relative pathogenicity of human genetic variants. Nature genetics, 46(3):310–315, 2014.

[43] Karthik A Jagadeesh, Aaron M Wenger, Mark J Berger, Harendra Guturu, Peter D Stenson, David N Cooper, Jonathan A Bernstein, and Gill Bejerano. M-cap eliminates a majority of variants of uncertain significance in clinical exomes at high sensitivity. Nature genetics, 48(12): 1581, 2016.

[44] Laksshman Sundaram, Hong Gao, Samskruthi Reddy Padigepati, Jeremy F McRae, Yanjun Li, Jack A Kosmicki, Nondas Fritzilas, Jörg Hakenberg, Anindita Dutta, John Shon, et al. Predicting the clinical impact of human mutation with deep neural networks. Nature genetics, 50(8):1161–1170, 2018.

[45] Haicang Zhang, Michelle S Xu, Wendy K Chung, and Yufeng Shen. Predicting functional effect of missense variants using graph attention neural networks. bioRxiv, 2021.

[46] Ngak Leng Sim, Prateek Kumar, Jing Hu, Steven Henikoff, Georg Schneider, and Pauline C. Ng. SIFT web server: Predicting effects of amino acid substitutions on proteins. Nucleic Acids Research, 40(W1):W452, jul 2012. ISSN 03051048. doi: 10.1093/nar/gks539.

[47] Adam Riesselman, Jung-Eun Shin, Aaron Kollasch, Conor McMahon, Elana Simon, Chris Sander, Aashish Manglik, Andrew Kruse, and Debora Marks. Accelerating Protein Design Using Autoregressive Generative Models. bioRxiv, page 757252, 2019. doi: 10.1101/757252. URL https://www.biorxiv.org/content/10.1101/757252v1.

[48] Brian L Hie, Kevin K Yang, and Peter S Kim. Evolutionary velocity with protein language models. bioRxiv, 2021.

[49] Henrik Nordberg, Michael Cantor, Serge Dusheyko, Susan Hua, Alexander Poliakov, Igor Shabalov, Tatyana Smirnova, Igor V Grigoriev, and Inna Dubchak. The genome portal of the department of energy joint genome institute: 2014 updates. Nucleic acids research, 42(D1): D26–D31, 2014.

[50] M. Steinegger, M. Mirdita, and J. Söding. Protein-level assembly increases protein sequence recovery from metagenomic samples manyfold. Nat Methods, 16(7):603–606, 07 2019.

[51] Alan S. Lapedes, Bertrand G. Giraud, LonChang Liu, and Gary D. Stormo. Correlated Mutations in Models of Protein Sequences: Phylogenetic and Structural Effects. Lecture Notes-Monograph Series, 33:236–256, 1999. doi: 10.2307/4356049. URL http://www.jstor.org/stable/4356049.

[52] John Thomas, Naren Ramakrishnan, and Chris Bailey-Kellogg. Graphical models of residue coupling in protein families, 4 2008. ISSN 15455963. URL https://pubmed.ncbi.nlm.nih.gov/18451428/.

[53] Martin Weigt, Robert A. White, Hendrik Szurmant, James A. Hoch, and Terence Hwa. Identification of direct residue contacts in protein-protein interaction by message passing. Proceedings of the National Academy of Sciences of the United States of America, 106(1):67–72, 1 2009. ISSN 00278424. doi: 10.1073/pnas.0805923106. URL https://www.pnas.org/content/106/1/67https://www.pnas.org/content/106/1/67.abstract.

[54] Sivaraman Balakrishnan, Hetunandan Kamisetty, Jaime G. Carbonell, Su-In Lee, and Christopher James Langmead. Learning generative models for protein fold families. Proteins: Structure, Function, and Bioinformatics, 79(4):1061–1078, 4 2011. ISSN 08873585. doi: 10.1002/prot.22934. URL http://doi.wiley.com/10.1002/prot.22934.

[55] David T. Jones, Daniel W. A. Buchan, Domenico Cozzetto, and Massimiliano Pontil. PSI-COV: precise structural contact prediction using sparse inverse covariance estimation on large multiple sequence alignments. Bioinformatics, 28(2):184–190, 1 2012. ISSN 1460-2059. doi: 10.1093/bioinformatics/btr638. URL https://academic.oup.com/bioinformatics/article-lookup/doi/10.1093/bioinformatics/btr638.

[56] Emily E Wrenbeck, Matthew S Faber, and Timothy A Whitehead. Deep sequencing methods for protein engineering and design. Current opinion in structural biology, 45:36–44, 2017.

[57] Justin R Klesmith, John-Paul Bacik, Ryszard Michalczyk, and Timothy A Whitehead. Comprehensive sequence-flux mapping of a levoglucosan utilization pathway in e. coli. ACS synthetic biology, 4(11):1235–1243, 2015.

[58] Hugh K Haddox, Adam S Dingens, Sarah K Hilton, Julie Overbaugh, and Jesse D Bloom. Mapping mutational effects along the evolutionary landscape of hiv envelope. Elife, 7:e34420, 2018.

[59] Philip A Romero, Tuan M Tran, and Adam R Abate. Dissecting enzyme function with microfluidic-based deep mutational scanning. Proceedings of the National Academy of Sciences, 112(23):7159–7164, 2015.

[60] Elad Firnberg, Jason W Labonte, Jeffrey J Gray, and Marc Ostermeier. A comprehensive, high-resolution map of a gene’s fitness landscape. Molecular biology and evolution, 31(6): 1581–1592, 2014.

[61] Zhifeng Deng, Wanzhi Huang, Erol Bakkalbasi, Nicholas G Brown, Carolyn J Adamski, Kacie Rice, Donna Muzny, Richard A Gibbs, and Timothy Palzkill. Deep sequencing of systematic combinatorial libraries reveals *β*-lactamase sequence constraints at high resolution. Journal of molecular biology, 424(3-4):150–167, 2012.

[62] Michael A Stiffler, Doeke R Hekstra, and Rama Ranganathan. Evolvability as a function of purifying selection in tem-1 *β*-lactamase. Cell, 160(5):882–892, 2015.

[63] Hervé Jacquier, André Birgy, Hervé Le Nagard, Yves Mechulam, Emmanuelle Schmitt, Jérémy Glodt, Beatrice Bercot, Emmanuelle Petit, Julie Poulain, Guilène Barnaud, et al. Capturing the mutational landscape of the beta-lactamase tem-1. Proceedings of the National Academy of Sciences, 110(32):13067–13072, 2013.

[64] Scott D Findlay and Lynne-Marie Postovit. Comprehensive characterization of transcript diversity at the human nodal locus. BioRxiv, page 254409, 2018.

[65] Richard N McLaughlin Jr, Frank J Poelwijk, Arjun Raman, Walraj S Gosal, and Rama Ranganathan. The spatial architecture of protein function and adaptation. Nature, 491(7422): 138–142, 2012.

[66] Jacob O Kitzman, Lea M Starita, Russell S Lo, Stanley Fields, and Jay Shendure. Massively parallel single-amino-acid mutagenesis. Nature methods, 12(3):203–206, 2015.

[67] Michael B Doud and Jesse D Bloom. Accurate measurement of the effects of all amino-acid mutations on influenza hemagglutinin. Viruses, 8(6):155, 2016.

[68] K Pokusaeva, C Johnson, B Luk, G7 Uribe, Y Fu, N Oezguen, RK Matsunami, M Lugo, A Major, Y Mori-Akiyama, et al. Gaba-producing bifidobacterium dentium modulates visceral sensitivity in the intestine. Neurogastroenterology & Motility, 29(1):e12904, 2017.

[69] Parul Mishra, Julia M Flynn, Tyler N Starr, and Daniel NA Bolon. Systematic mutant analyses elucidate general and client-specific aspects of hsp90 function. Cell reports, 15(3):588–598, 2016.

[70] Eric D Kelsic, Hattie Chung, Niv Cohen, Jimin Park, Harris H Wang, and Roy Kishony. Rna structural determinants of optimal codons revealed by mage-seq. Cell systems, 3(6):563–571, 2016.

[71] Alexandre Melnikov, Peter Rogov, Li Wang, Andreas Gnirke, and Tarjei S Mikkelsen. Comprehensive mutational scanning of a kinase in vivo reveals substrate-dependent fitness landscapes. Nucleic acids research, 42(14):e112–e112, 2014.

[72] Lisa Brenan, Aleksandr Andreev, Ofir Cohen, Sasha Pantel, Atanas Kamburov, Davide Cacchiarelli, Nicole S Persky, Cong Zhu, Mukta Bagul, Eva M Goetz, et al. Phenotypic characterization of a comprehensive set of mapk1/erk2 missense mutants. Cell reports, 17(4):1171–1183, 2016.

[73] Liat Rockah-Shmuel, Ágnes Tóth-Petróczy, and Dan S Tawfik. Systematic mapping of protein mutational space by prolonged drift reveals the deleterious effects of seemingly neutral mutations. PLoS computational biology, 11(8):e1004421, 2015.

[74] Nicholas C Wu, C Anders Olson, Yushen Du, Shuai Le, Kevin Tran, Roland Remenyi, Danyang Gong, Laith Q Al-Mawsawi, Hangfei Qi, Ting-Ting Wu, et al. Functional constraint profiling of a viral protein reveals discordance of evolutionary conservation and functionality. PLoS genetics, 11(7):e1005310, 2015.

[75] Christopher D Aakre, Julien Herrou, Tuyen N Phung, Barrett S Perchuk, Sean Crosson, and Michael T Laub. Evolving new protein-protein interaction specificity through promiscuous intermediates. Cell, 163(3):594–606, 2015.

[76] Hangfei Qi, C Anders Olson, Nicholas C Wu, Ruian Ke, Claude Loverdo, Virginia Chu, Shawna Truong, Roland Remenyi, Zugen Chen, Yushen Du, et al. A quantitative high-resolution genetic profile rapidly identifies sequence determinants of hepatitis c viral fitness and drug sensitivity. PLoS pathogens, 10(4):e1004064, 2014.

[77] Kenneth A Matreyek, Lea M Starita, Jason J Stephany, Beth Martin, Melissa A Chiasson, Vanessa E Gray, Martin Kircher, Arineh Khechaduri, Jennifer N Dines, Ronald J Hause, et al. Multiplex assessment of protein variant abundance by massively parallel sequencing. Nature genetics, 50(6):874–882, 2018.

[78] Pradeep Bandaru, Neel H Shah, Moitrayee Bhattacharyya, John P Barton, Yasushi Kondo, Joshua C Cofsky, Christine L Gee, Arup K Chakraborty, Tanja Kortemme, Rama Ranganathan, et al. Deconstruction of the ras switching cycle through saturation mutagenesis. Elife, 6:e27810, 2017.

[79] Benjamin P Roscoe, Kelly M Thayer, Konstantin B Zeldovich, David Fushman, and Daniel NA Bolon. Analyses of the effects of all ubiquitin point mutants on yeast growth rate. Journal of molecular biology, 425(8):1363–1377, 2013.

[80] Benjamin P Roscoe and Daniel NA Bolon. Systematic exploration of ubiquitin sequence, e1 activation efficiency, and experimental fitness in yeast. Journal of molecular biology, 426(15): 2854–2870, 2014.

[81] David Mavor, Kyle Barlow, Samuel Thompson, Benjamin A Barad, Alain R Bonny, Clinton L Cario, Garrett Gaskins, Zairan Liu, Laura Deming, Seth D Axen, et al. Determination of ubiquitin fitness landscapes under different chemical stresses in a classroom setting. Elife, 5: e15802, 2016.

[82] Yvonne H Chan, Sergey V Venev, Konstantin B Zeldovich, and C Robert Matthews. Correlation of fitness landscapes from three orthologous tim barrels originates from sequence and structure constraints. Nature communications, 8(1):1–12, 2017.

[83] Daniel Melamed, David L Young, Caitlin E Gamble, Christina R Miller, and Stanley Fields. Deep mutational scanning of an rrm domain of the saccharomyces cerevisiae poly (a)-binding protein. Rna, 19(11):1537–1551, 2013.

[84] Lea M Starita, Jonathan N Pruneda, Russell S Lo, Douglas M Fowler, Helen J Kim, Joseph B Hiatt, Jay Shendure, Peter S Brzovic, Stanley Fields, and Rachel E Klevit. Activity-enhancing mutations in an e3 ubiquitin ligase identified by high-throughput mutagenesis. Proceedings of the National Academy of Sciences, 110(14):E1263–E1272, 2013.

[85] Carlos L Araya, Douglas M Fowler, Wentao Chen, Ike Muniez, Jeffery W Kelly, and Stanley Fields. A fundamental protein property, thermodynamic stability, revealed solely from large-scale measurements of protein function. Proceedings of the National Academy of Sciences, 109 (42):16858–16863, 2012.

[86] Julian Salazar, Davis Liang, Toan Q. Nguyen, and Katrin Kirchhoff. Pseudolikelihood reranking with masked language models. CoRR, abs/1910.14659, 2019. URL http://arxiv.org/abs/1910.14659.

[87] Kevin K. Yang, Zachary Wu, Claire N. Bedbrook, and Frances H. Arnold. Learned protein embeddings for machine learning. Bioinformatics, 34(15):2642–2648, 8 2018. ISSN 14602059. doi: 10.1093/bioinformatics/bty178.

[88] Andrew Leaver-Fay, Matthew J O’meara, Mike Tyka, Ron Jacak, Yifan Song, Elizabeth H Kellogg, James Thompson, Ian W Davis, Roland A Pache, Sergey Lyskov, et al. Scientific benchmarks for guiding macromolecular energy function improvement. Methods in enzymology, 523:109–143, 2013.

[89] Milot Mirdita, Lars Von Den Driesch, Clovis Galiez, Maria J. Martin, Johannes Soding, and Martin Steinegger. Uniclust databases of clustered and deeply annotated protein sequences and alignments. Nucleic Acids Research, 45(D1):D170–D176, 1 2017. ISSN 13624962. doi: 10.1093/nar/gkw1081.

[90] Lukas Neumann, Andrew Zisserman, and Andrea Vedaldi. Relaxed Softmax: Efficient Confidence Auto-Calibration for Safe Pedestrian Detection. Technical report, oct 2018.

[91] Jeremy Nixon, Mike Dusenberry, Ghassen Jerfel, Timothy Nguyen, Jeremiah Liu, Linchuan Zhang, and Dustin Tran. Measuring Calibration in Deep Learning. apr 2019. URL http://arxiv.org/abs/1904.01685.

[92] L. Steven Johnson, Sean R. Eddy, and Elon Portugaly. Hidden Markov model speed heuristic and iterative HMM search procedure. BMC Bioinformatics, 11(1):431, 8 2010. ISSN 14712105. doi: 10.1186/1471-2105-11-431. URL https://bmcbioinformatics.biomedcentral.com/articles/10.1186/1471-2105-11-431.

[93] Didier Nurizzo, Steven C Shewry, Michael H Perlin, Scott A Brown, Jaydev N Dholakia, Roy L Fuchs, Taru Deva, Edward N Baker, and Clyde A Smith. The crystal structure of aminoglycoside-3-phosphotransferase-iia, an enzyme responsible for antibiotic resistance. Journal of molecular biology, 327(2):491–506, 2003.

